# Human pluripotent stem cells-derived inner ear organoids recapitulate otic development *in vitro*

**DOI:** 10.1101/2023.04.11.536448

**Authors:** Daniela Doda, Sara Alonso Jimenez, Hubert Rehrauer, Jose F. Carreño, Victoria Valsamides, Stefano Di Santo, Hans Ruedi Widmer, Albert Edge, Heiko Locher, Wouter van der Valk, Jingyuan Zhang, Karl R. Koehler, Marta Roccio

## Abstract

Our molecular understanding of the early stages of human inner ear development has been limited by the difficulty in accessing fetal samples at early gestational stages. As an alternative, previous studies have shown that inner ear morphogenesis can be partially recapitulated using induced pluripotent stem cells (iPSCs) directed to differentiate into Inner Ear Organoids (IEOs). Once validated and benchmarked, these systems could represent unique tools to complement and refine our understanding of human otic differentiation and model developmental defects. Here, we provide the first direct comparisons of the early human embryonic otocyst and human iPSC-derived IEOs. We use multiplexed immunostaining, and single-cell RNA sequencing to characterize IEOs at three key developmental steps, providing a new and unique signature of *in vitro* derived otic -placode, -epithelium, -neuroblasts, and -sensory epithelia. In parallel, we evaluate the expression and localization of critical markers at these equivalent stages in human embryos. We show that the placode derived *in vitro* (days 8-12) has similar marker expression to the developing otic placode of Carnegie Stage (CS) 11 embryos and subsequently (days 20-40) this gives rise to otic epithelia and neuroblasts comparable to the CS13 embryonic stage. Differentiation of sensory epithelia, including supporting cells and hair cells starts *in vitro* at days 50-60 of culture. The maturity of these cells is equivalent to vestibular sensory epithelia at week 10 or cochlear tissue at week 12 of development, before functional onset. Together, our data indicate that the current state-of-the-art protocol enables the specification of *bona fide* otic tissue, supporting the further application of IEOs to inform inner ear biology and disease.

## Introduction

The inner ear contains six distinct epithelial sensory patches that detect head movement- and sound-induced fluid vibration and act as peripheral receptors for the senses of balance and hearing. Sensory epithelia comprise two cell types: supporting cells and hair cells. The latter function as mechanoreceptors. Mechanical deflection of stereocilia bundles in the apical domain of hair cells results in cellular depolarization and neurotransmitter release on the basal side. This in turn, activates the afferent fibers of the cochlear or vestibular neurons in direct synaptic contact with them, which subsequently relay to the upper auditory centers. Sensory and non-sensory epithelia, as well as the cochleovestibular neurons have a common developmental origin in the otic vesicle (OV) or otocyst (Alsina and Whitfield, 2017; Groves and Fekete, 2012). The OV is derived from the invagination of the otic placode (OP) into the underlying mesenchyme. The OV forms around the fourth week (W4) of human embryo development (O’Rahilly, 1963), and in mice around embryonic days 8.5-9 (E8.5-E9) (Wu and Kelley, 2012).

The OP is one of the posterior placodes (Saint-Jeannet and Moody, 2014). Cranial placodes are specified from a transient pre-placodal ectoderm (PPE) region that develops at the neural border, between the neural plate and non-neural ectoderm (Litsiou et al., 2005; Martin and Groves, 2006; Saint-Jeannet and Moody, 2014; Steventon et al., 2014). This is further patterned along the anterior-posterior axis of the developing head and subdivides into distinct placodes giving rise to the lens, the nasal/olfactory, the trigeminal, the adenohypophyseal, the otic and the epibranchial placodes. The OP is characterized by the expression of genes such as *PAX8 and PAX2*, which are also required for inner ear morphogenesis (Bouchard et al., 2010). Upon invagination and detachment from the surface ectoderm, complex morphogenetic events lead to the generation of the inner ear membranous labyrinth and its’ specialized sensory organs (Groves and Fekete, 2012; Wu and Kelley, 2012). Morphogens secreted by the hindbrain, the mesenchyme and the floor plate, generate the coordinates for the anterior-posterior, the lateral-medial and the dorsal-ventral axis in the OV and result in the specification of sensory and non-sensory regions, as well as cochlear or vestibular fate (Bok et al., 2007; Durruthy-Durruthy et al., 2014; Riccomagno et al., 2005; Wu and Kelley, 2012).

The ventral portion of the OV contains SOX2-expressing progenitors that form the so-called pro-neurosensory domain (Kiernan et al., 2005; Puligilla et al., 2010; Steevens et al., 2017). This area gives rise to both the sensory epithelia and the neural components of the inner ear. Neuroblasts delaminate from the pro-neurosensory region (Evsen et al., 2013 ; Puligilla et al., 2010; Radde-Gallwitz et al., 2004; Steevens et al., 2017) and coalesce to form the cochleovestibular ganglion (CVG) which will further innervate the sensory patches. In mouse development this occurs around E9.5-E10 (Appler and Goodrich, 2011). The prosensory region within the otocyst further segregates into distinct prosensory domains which give rise to the sensory epithelia in the ampullae of the semicircular canals, to the maculae of the utricle and saccule and to the sensory epithelium in the cochlear duct (Gu et al., 2016). Migrating cranial neural crest (CNC) infiltrates the OV and surrounding otic mesenchyme and contributes to the population of glial/Schwann cells in the cochlear and vestibular ganglia (Mendez-Maldonado et al., 2020; Renauld et al., 2022) and to the intermediate cell layer in the stria vascularis (Adameyko et al., 2012; Cable et al., 1992; Locher et al., 2015; Renauld et al., 2022; Steel and Barkway, 1989). CNC-derived mesenchyme instead, will give rise to the ossicles of the middle ear (Martik and Bronner, 2021).

Our knowledge of these early developmental events is almost exclusively based on studies performed in model organisms such as the mouse and the chick; It is unclear how similarly these processes play out in humans. We and others have shown that the appearance of the first hair cells in the vestibular organs occurs around weeks 8-10 of human development. In the cochlea instead, these develop a few days later, with a basal to apical gradient starting at week 11 (Lavigne-Rebillard and Pujol, 1986; Locher et al., 2014; Locher et al., 2013; Pechriggl et al., 2015; Pujol and Lavigne-Rebillard, 1985; Roccio et al., 2018; Sans and Dechesne, 1985). By week 20, hair bundles can be observed (Lavigne-Rebillard and Pujol, 1987; Lavigne-Rebillard and Pujol, 1988) and *in-utero* hearing starts during the third trimester. Using multiplexed qPCR we previously provided the first molecular characterization of the developing cochlear duct, utricular macula and spiral ganglion at weeks 8-12 of fetal development (Roccio et al., 2018). More recent studies exploiting single-cell RNA sequencing have provided additional insights into the transcriptional profile of the human inner ear components (Shengyang Yu et al.; van der Valk et al., 2022). Earlier stages of human inner ear development are instead scarcely characterized. While anatomical studies based on CT scans/MRI (Yasuda et al., 2007) or tissue sectioning followed by 3D reconstruction (de Bakker et al., 2016) and embryology atlases (https://www.3dembryoatlas.com; https://www.ehd.org/virtual-human-embryo/) give us a picture of morphological events during early otocyst/inner ear development, the molecular characterization of this process is limited. Major experimental, logistic, and ethical limitations remain, which hamper the availability and accessibility to human fetal/embryonic inner ear tissue.

State-of-the-art inner ear organoids (IEO) protocols based on stem cell differentiation in 3D culture recapitulate many features of inner ear morphogenesis *in vitro* (Koehler et al., 2017; Munnamalai and Fekete, 2017; van der Valk et al., 2021). Pluripotent stem cells are guided in a stepwise fashion towards the otic lineage, eventually leading to multicellular aggregates bearing sensory and non-sensory inner ear epithelia and otic-like neurons (Koehler et al., 2017; Nie and Hashino, 2020). IEOs are a powerful tool to refine our understanding of the molecular signals leading to inner ear cell type specification. Benchmarking these models to the primary human tissue would provide a robust base for their future use to study organ development and model diseases. In addition, reliable models of the human inner ear could allow for testing novel therapeutic strategies for sensorineural hearing loss and vestibular disorders (Nist-Lund et al., 2022; Roccio and Edge, 2019).

Here, we have compared for the first time a set of human embryos from Carnegie stage 11 to 13 (week 4-5 of embryonic development), to *in vitro* stages of IEO differentiation. We observed the expression of shared genes and protein markers between CS11 human embryos and early-stage IEOs (day 8-12 *in vitro*). Otic vesicle-like structures and neuroblasts derived *in vitro* at days 20-40 showed remarkable similarity to CS13 embryos. Finally, sensory vesicles matured to days 50-60 were comparable to weeks 10-12 inner ear fetal specimens. Our findings indicate that IEOs faithfully recapitulate otic developmental steps, by generating the progenitors of the inner ear sensory and non-sensory lineages and provide a solid foundation for future studies of human inner ear biology.

## Material and Methods

### Human fetal sample collection

Human samples were collected in two different medical centres. Procurement and procedures were performed with full approval by the respective Ethics Committee. All experiments were performed in accordance with guidelines enunciated in the Declaration of Helsinki (DoH) and the guide lines of the international society for stem cell research (ISSCR).

Samples: E1026, E1027, E1037, E852, E1291, E1203; E1201 were collected as previously described (Roccio et al., 2018). Signed informed consent of the donors for procurement of the aborted embryos/fetuses and for use of tissues in research was obtained. Procedures were performed with full approval by the Ethics Committee of the Medical Faculty of the University of Bern and the Ethics Committee of the State Bern, Switzerland (Gesundheits-und Fürsorgedirektion des Kantons Bern, Kantonale Ethikkommission für die Forschung (Project ID: 2016–00033/KEK-Nr. 181/ 07). Tissues were collected the same day, as early as possible after the procedure, otherwise discarded. Samples were then anonymised. Staging was performed by comparing tissue morphology to atlases (de Bakker et al., 2016) or as previously described (Evtouchenko L, 1996; Roccio et al., 2018). The sample EF1 was collected at the Leiden University Medical Center (LUMC) according to the Dutch legislation (Fetal Tissue Act, 2001), under the protocol number B19.070 approved by the Medical Research Ethics Committee of LUMC as well as written informed consent of the donor following the Guidelines on the Provision of Fetal Tissue set by the Dutch Ministry of Health, Welfare, and Sport (revised version, 2018).The inner ear tissue was collected after elective termination of pregnancy by vacuum aspiration. Embryonic or fetal age was determined by obstetric ultrasonography prior to termination, with a standard error of 2 days. All references to human developmental weeks refer to fetal age.

After collection, samples were fixed overnight in paraformaldehyde (PFA), in PBS pH 7.4 (11762.00250, *Morphisto)* at 4°C and subsequently placed in PBS (14190169, Thermo Fisher / Life Technologies).

### iPSC lines and culture

Cell lines (hiPSC SOX2-GFP) and (hiPSC LMNB1-RFP), derived from the parental WTC-11 hiPSC were obtained from the Coriell Institute and generated at the Allen Institute for Cell Science at passage 29 frozen in mTeSR™ Plus medium (STEMCELL Technologies). The PAX2-EGFP-iPSC line was generated starting from BJ2 iPSC lines at the Harvard Stem Cell Institute core facility. BJ fibroblasts (ATCC) were reprogrammed to generate iPSC. Knock-in was performed using CRISPR-Cas9 (the generation of the line is part of a manuscript in preparation. Details available on request). Clone C2 with bi-allelic insertion was used for the experiments given the brighter GFP fluorescence. Cells were obtained at p34, frozen in mTESR1 medium.

Cells were thawed according to the provider’s protocol and the medium was gradually changed to Essential 8 Flex Medium (E8f) (A2858501 Thermo Fisher / Life Technologies) supplemented with 100 μg/ml Normocin (4069-ant-nr-1, InvivoGen), on recombinant human Vitronectin (A14700, Thermo Fisher / Life Technologies)-coated 6-well plates. 1 ml PBS containing 0.5 mg/ml Vitronectin was used to coat each well for at least 1 hour at 37°C prior to cell seeding. Cells were passaged at circa 70-80% confluency or every 4 or 5 days using Accutase (7002353, Thermo Fisher / Life Technologies). Briefly, the medium was removed, cells were rinsed with PBS and incubated for 4-5 min with 500μl Accutase at 37°C. 1ml of the medium was subsequently added to the plates for cell dissociation and cells were quickly centrifuged (300g, 5min). The cell pellet was resuspended in E8f with 10μM ROCKi (Y-27632, ALX-270-333-M005, EnzoLifeSciences) and replated. The subsequent day the medium was changed to remove the ROCKi, and subsequently changed as according to medium manufacturer’s instructions. iPSC lines were analyzed for expression of pluripotency markers SOX2, OCT4, and SSEA-1, TRA1-60 and reporter expression. Cells obtained from the Coriell institute at passage 29 were used between passages 30-35. Cells obtained from the HSCI core facility at passage 34 were used between passages 35-40.

### Inner Ear Organoid differentiation

iPSC differentiation to inner ear organoid followed previously described protocols (Koehler et al., 2017; Nie and Hashino, 2020) with some modification. Briefly, 70-80% confluent iPSC cultures were dissociated to a single cell suspension in E8f supplemented with 20μM ROCKi. 3500 cells per well, were aggregated in round bottom low-binding 96 well plates (174929, Thermo Fisher / Life Technologies) by centrifugation (100g, 6 min) in 100 μl E8f medium with 20μM ROCKi. The following day, the addition of 100 μl E8f was used to dilute the ROCKi to 10μM.

On day 2 of culture (day 0 differentiation) the organoids, *circa* 500μm in diameter, were collected, rinsed twice in E6 medium (A1516401, Thermo Fisher / Life Technologies) and subsequently transferred to differentiation medium “E6/SB/F/B4” (see below) with 1 organoid/100μl differentiation medium/well in new round bottom low-binding plates.

Differentiation medium E6/SB/F/B4 consisted of E6 supplemented with 2% growth factors-reduced Matrigel (354230, Corning), SB-431542 at 10μM (CAY13031, Cayman Chemical) 4ng/ml FGF2 (SRP3043, Merk/Sigma-Aldrich) and BMP-4 (314-BP-010 Biotechne/R&D Systems) from 0.5-10ng/ml. The optimal concentration for placode induction with the cell line used ranged from 1.5-2.5 ng/ml. On days 3-4 of differentiation, FGF2 and LDN193189 (CAY19396, Cayman Chemical) were added to the medium at the final concentration of 50ng/ml and 200nM respectively. On day 8, the medium was supplemented with 3μM CHIR99021 (361571 Avantor/WVR) to promote otic fate. CHIR99021 was further supplemented on day 10. On day 12 of *in vitro* culture, the organoids were transferred to Organoid Maturation Medium (OMM) consisting of 1:1 Neurobasal (21103049, Thermo Fisher / Life Technologies) : Advanced-DMEMF12 (12634028, Thermo Fisher / Life Technologies), supplemented with 0.5X B27 (12587010, Thermo Fisher / Life Technologies); 0.5X N2 (17502048, Thermo Fisher / Life Technologies), 1XGlutamax (35050061,Thermo Fisher / Life Technologies), 1X beta mercapto ethanol (21985023, Gibco/Thermo Fisher / Life Technologies) and CHIR99021 3μM.

Organoids were transfered 1 organoid per well in 24-well plates (Non-treated) (144530 Thermo Fisher / Life Technologies) pre-coated as follows: 500μl anti-adherent solution (STEMCELL Technologies) was added to each well for 5 minutes at room temperature. The surfactant was then removed by aspiration, and wells were washed twice with 1ml PBS and dried for 5 minutes prior to use.

The medium was refreshed to provide fresh CHIR on day 15 of differentiation. From day 18 onwards, half of the OMM medium was changed every 2 days and refreshed with OMM without Matrigel. Samples were collected at selected time points and either fixed for characterization or dissociated to single-cells for flow cytometry/scRNA seq.

### Organoids and embryos characterization

#### Cryosections

Organoids were collected using wide-opening pipet tips at different time points and fixed in 4% PFA, in PBS pH 7.4 for 15 minutes at room temperature, subsequently rinsed in PBS and incubated for 24 hours in 18% sucrose (S0389, Sigma Aldrich) solution at 4°C and subsequently, transferred for another 24 hours in 30% sucrose in PBS. Organoids were then transferred to OCT medium (361603E, Avantor/*VWR*) in cryomolds and then frozen on a dry-ice bed. Cryoblocks were incubated at last 1 day at −80°C prior to sectioning. The same procedure was used for human samples (CS11-13). For the W12 cochleae, these were decalcified with 0.5M EDTA (15575020 Thermo Fisher / Life Technologies) in PBS for 30 days prior to preparation for cutting as cryosections.

Cryoblocks were sectioned at a thickness of 12μm with a Cryostat (Leica CM3050 S) and sections were collected on Superfrost Plus Adhesion microscope slides (J1800AMNZ, Epredia). For experiment where organoids were exposed to different conditions during the differentiation phase, all conditions were collected on the same glass slide for further comparison with antibody-based staining.

#### Paraffin embedding end sectioning

Samples embedding in paraffin was performed as follows: After fixation samples dehydrated in an increasing ethanol series (70%–99%; 84050068.2500, Boom), cleared in xylene (534056, Honeywell), and embedded in paraffin wax (2079, Klinipath). Sections of 4-5 μm were cut using a rotary microtome (HM355S, Thermo Scientific). Sections were deparaffinized in xylene and rehydrated in a descending series of ethanol (96%–50%) and several rinses in deionized water. Before immunostaining, antigen retrieval was performed in 10 mM sodium citrate buffer (pH 6.0; S1804-500G, Sigma-Aldrich) for 12 min at in a microwave at *circa* 97°C

#### Imaging and image analysis

Samples were permeabilized with PBS containing 0.1% Triton-X100 (X100, Sigma Aldrich) and subsequently incubated with blocking buffer (BB) containing 2% BSA (A2153 Sigma Aldrich) and 0.01% Triton-X100 in PBS for 2 hours at room temperature. Finally, each slide was incubated with 250μl of antibody containing solution in BB. The used antibodies are indicated in **Table 1**. After primary antibody incubation (overnight at 4°C or 2 hours at room temperature), samples were rinsed three times for 10 minutes with PBS and incubated with Alexa-Fluor conjugated antibodies (**Table 1**) (overnight at 4°C or 2hs at room temperature). Finally, slides were washed with PBS and mounted using glycerol mounting medium with DABCO (INTFP-WU1420, Interchim)

### Image acquisition and quantification

Widefield images were acquired with a Zeiss Axio-observer Z1 microscope with a motorized stage. Filter sets: DAPI (ex: BP390/22-25 em: BP 455/50); GFP (ex:BP484/25 em: BP525/50) DsRed (ex:BM553/18 em: BM607/70) CY5 (ex: BP: 631/22 em: BP 690/50). For organoid assessment, different experimental conditions (or time points) were positioned on the same glass slides (see also **Figure S2B**). Tile scans were acquired for comparative assessment of marker expression. Single images of organoids/sections were then acquired using a 10x (EC Plan Neufluar) or 20x (LD Plan Neufluar) objectives.

Areas of interest were marked and re-imaged using a laser-scanning confocal microscope. Confocal images were acquired using a Leica SP8 microscope either in 10x (HC PL Fluotar) air objective; a 20x (PL APO CS) air objective or a 63x (HC PL APO CS2 NA 1.4) oil objective. Sequential imaging was used to acquire different channels.

To assess marker-positive areas, image files were opened using FIJI (Schindelin et al., 2012). The contour of the organoids was drawn manually on the DAPI channel and used to quantify the organoid area. Within this ROI, a fluorescent threshold was placed-on the channel of the marker of interest-to identify signal-positive areas. Alternatively, the vesicle contour was manually and the lumen volume subtracted. For organoids containing multiple vesicles, the sum of all areas of each vesicle was calculated and expressed as a ratio to the area of the organoid section. 2-3 images of the same organoids and 3-4 organoids per condition were analyzed.

If confocal images with multiple stacks were acquired, images are shown as a 3D projection stack.

### Organoid dissociation for scRNAseq

Organoids were collected at the defined time points in a 15 ml falcon tube and let sediment by gravity. After removing the culture medium, organoids were incubated with Accutase (1X) and DNAse1 0.5mg/ml (10104159001, sigma Aldrich) for 25-30 min at 37°C in a water bath, in 1 ml dissociation solution per plate (at d8) or per ½ plate (>d18). Every 5 minutes, organoids were triturated mechanically by pipetting with p1000 pipet tips 10 times. After 25-30 minutes, cells were washed with medium containing 10μM ROCKi and centrifuged for 5 minutes at 300g. The pellet was then resuspended in PBS containing 2% BSA and 10μM ROCKi and passed through a cell strainer with 40μm pore size (352235, Corning) to eliminate clumps. Cells were then further centrifuged and resuspended in 550μl PBS containing 2% BSA and 10μM ROCKi on ice.

Single-cell isolated from day 8 organoids (200 unselected organoids) were incubated for 5 minutes at room temperature with Calcein-AM (C3099, Life Technologies) and subsequently sorted on a FACS Aria (BD) to exclude debris, doublets, and isolate single positive (live) cells (these represented 96% of the cell suspension). Single cells from day 26 organoids (140 unselected organoids) were incubated on ice for 45 min with 40μl of an EPCAM-Pe-Cy7 antibody (25-9326 eBioscience/ThermoFisher Scientific) (0.36μg antibody in 500μl buffer), after washing, cells were sorted on a FACS Aria (BD) to purify Pe-Cy7+ cells. Debris and doublets were also excluded. Day 60 organoids (48 “unselected” organoids experiment 1) or (8 “selected” organoids experiment 2) were triturated as above and incubated for 30 min with 10μl (0.12ug) EPCAM-Pe-Cy7 antibody /1×10^6^ cells in 500μl buffer. After washing, cells were sorted on a FACS Aria (BD) to purify Pe-CY7+ cells. Unstained controls were used to define the gating strategy. The selection of organoids for experiment 2 was performed by live imaging of GFP fluorescence (SOX2-GFP) and brightfield imaging to visualize vesicles. Only organoids with visible vesicles were used. An example of such organoids is shown in **Fig. S8C**.

After sorting, cells were centrifuged for 10 minutes at 300g to remove FACS fluids and resuspended in a 0.04% BSA in PBS buffer at the concentration of 1000 cells/μl and placed on ice prior to loading on the 10X Chromium according to the manufacturer’s instruction. Sorting was performed at the Flowcytometry Facility of the University of Zurich.

### Single Cell RNA Sequencing and data analysis

The quality and concentration of the single-cell preparations were evaluated using a hemocytometer in a Leica DM IL LED microscope and adjusted to 400 cells/µl. 17,200 cells per sample were loaded into the 10X Chromium controller and library preparation was performed according to the manufacturer’s indications (Chromium Next GEM Single Cell 3ʹ Reagent Kits v3.1 protocol). The resulting libraries were sequenced in an Illumina NovaSeq sequencer according to 10X Genomics recommendations (paired-end reads, R1=28, i7=10, i5=10, R2=90) to a depth of around 40,000 reads per cell. Single-cell RNA-seq reads were processed with „cellranger count“ v6.1.2. to generate the single-cell gene expression matrix. Subsequently, we filtered doublet cells using the R/Bioconductor package „scDblFinder“ (Germain et al., 2021), and we removed low-quality cells using the quality metrics computed by the package „scater“ (McCarthy DJ, 2017). The remaining cells were analysed using the „Seurat“ package (Hao et al., 2021). The workflow included normalization, dimension reduction and clustering, as well as the identification of marker genes for clusters and differentially expressed genes.

Data sets at day 60 of differentiation obtained from two independent experiments were integrated using Seurat’s “IntegrateData” as described in the available online vignette (https://satijalab.org/seurat/articles/integration_introduction.html)(Hao et al., 2021). Selection of the population of interest (OEP and HC) was performed based on marker gene expression (https://satijalab.org/seurat/reference/addmodulescore). Modules including OEP genes and HC genes (**Table 2**) were used to select and further re-cluster these cell populations.

Data processing and analysis were performed using the Sushi platform (Hatakeyama et al., 2016). Data are visualized either as Uniform Manifold Approximation and Projection (UMAP) plots or as violin plots of normalized data.

Sequencing and bioinformatic analysis were performed at the Functional Genomics Center Zurich.

## Results

### Characterization of the developing human otocyst

The invagination of the otic placode into the head mesenchyme is a critical event leading to the formation of the otic vesicle. This occurs around day 20 of human development (Carnegie Stage (CS) 10-11) (de Bakker et al., 2016). Remarkably, two of the earliest fetal samples we collected (staged CS11, *circa* 24-25 days), provided a snapshot of otic vesicle (OV) formation and displayed otic pits in the process of pinching off from the surface ectoderm. We immunostained the CS11 specimens and later samples at CS12 (26-30 days) and CS13 (28-32 days) with markers of OV development (SOX2, PAX2, PAX8, SOX10), neural epithelium and new-born neurons (PAX6, TUBB3, NEUROD1) and neural crest (SOX10) to provide an initial molecular characterization of these samples (**Fig. 1, Fig. 2 and Fig. S1**).

**Figure 1:**
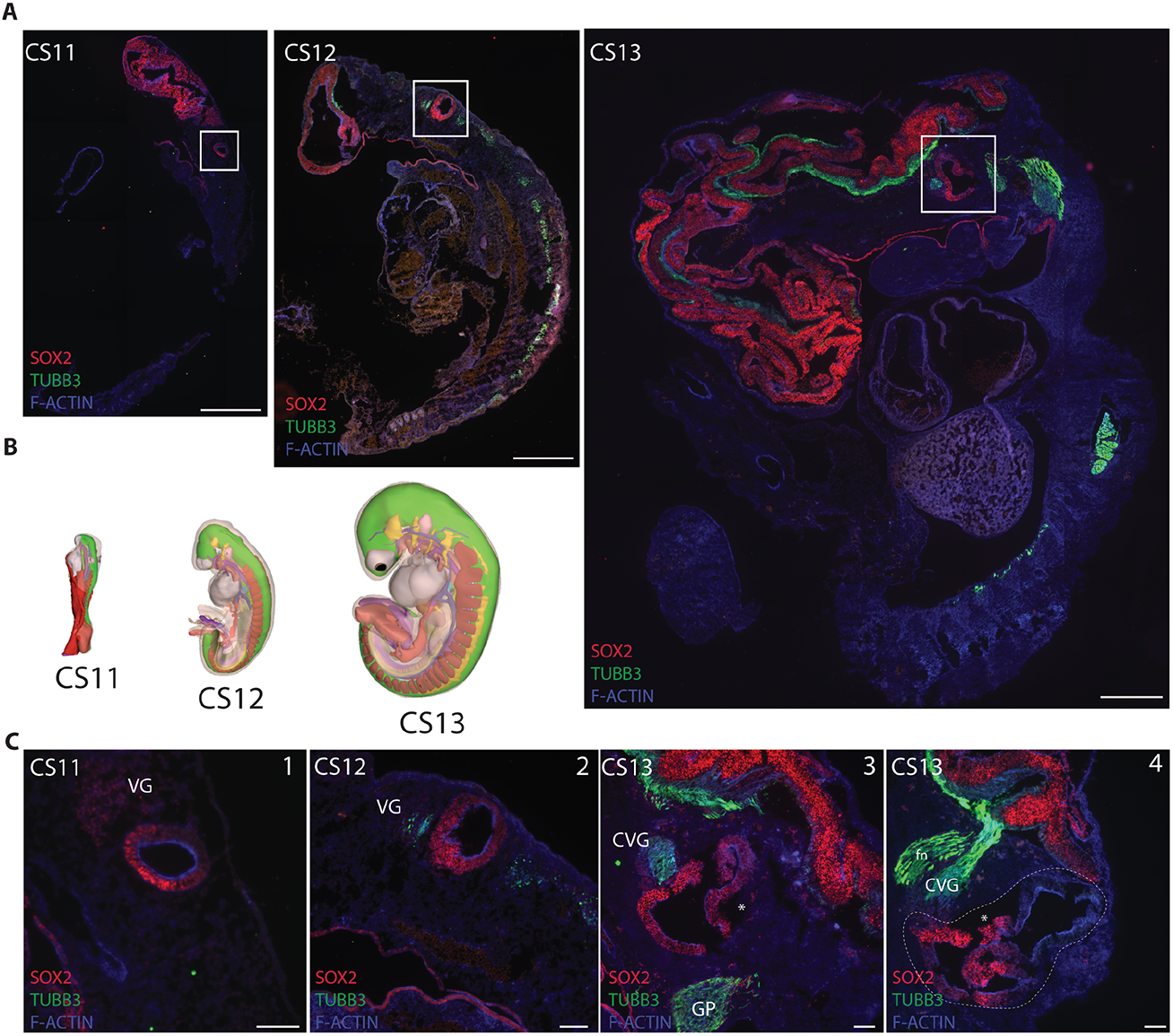
Otic vesicle development in human embryos. **A-** Human fetal samples stages CS11 (E1026), CS12 (E1027) and CS13 (E1037) immunolabelled to detect the developing otocysts (white box) with SOX2; TUBB3 and F-Actin (Phalloidin). Scale bar 1mm. **B-** 3D reconstructions embryos at CS11 (day 23-26), CS12 (day 26-30) and CS13 (day 28-32) adapted from the human development 3D atlas (https://www.3dembryoatlas.com). **C-** Higher magnification of the otocyst and the developing CVG ganglion stained with SOX2 and TUBB3 antibodies and F-Actin (Phalloidin). GP: Glossopharyngeal ganglion. Panels 3, and 4: sections at different lateral-medial levels of the same sample (E1037). fn-Putative branch of the facial nerve. Scale bar 100μm. *\* potential cutting artifact or tissue damage*

**Figure 2:**
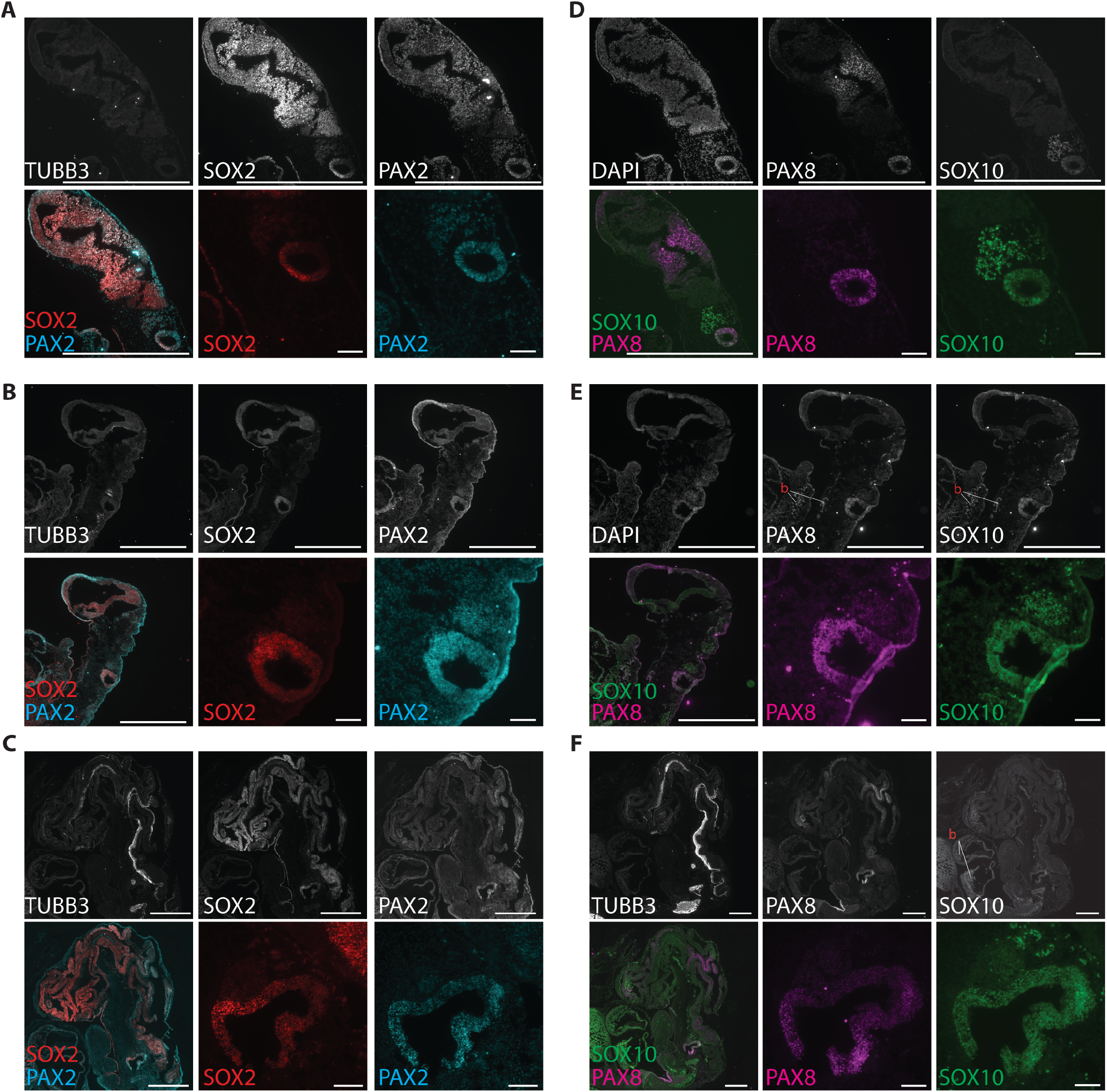
Characterization of otic vesicle development in human embryos. Human fetal samples stages CS11 (E1026) (**A** and **D**), CS12 (E1027) (**B** and **E**), CS13 (E1037) (**C** and **F**) immunolabelled to detect the developing otocysts. SOX2; PAX2 and TUBB3 staining (left). SOX10; PAX8, TUBB3 staining (right). A tile scan of the head region (scale bar 1mm) and a detail of the otocyst (scale bar 100μm) are shown for each panel. b: *Autofluorescence blood cells*

During this early developmental window, SOX2 expression is detectable in the developing CNS, as well as in the otocyst. In the latter, SOX2 staining appears predominantly localized in anteroventral portion (**Fig. 1C, Fig. 2A-C and Fig. S1B**). This is particularly obvious in sections at CS13, where the dorsal region of the otocyst/developing inner ear becomes visible (**Fig. 1C** panel 4), and it is in agreement with the pattern of expression in the murine otocyst (Gu et al., 2016; Steevens et al., 2017). βIII Tubulin (TUBB3) is undetectable in CS11 samples and starts to appear at CS12 (**Fig. 1 and Fig. S1E**). Here, TUBB3-positive cells appear in the region of the developing cochleovestibular ganglion (CVG). TUBB3 is moreover already expressed in other cranial ganglia, dorsal root ganglia (DRG), as well as in the developing brain (**Fig. S1E**). By CS13, CVG neurons project to the brainstem (**Fig. 1C** panel 4).

PAX2 expression is visible in the OV at CS11 (**Fig. 2A**). The staining becomes however more selective in the otocyst by CS13 (**Fig. 2C**). PAX2 can also be detected in different areas of the developing brain at all stages (**Fig. 2A-C**). PAX8 instead, is expressed in the otocyst from CS11 to CS13 and it is also present in the mid-hindbrain boundary region as previously reported in mice (Bouchard et al., 2010) (**Fig. 2D-F**). In addition, in other cranial ganglia and in the optic vesicle (**Fig. S1H**). Immunolabeling for SOX10 shows that at early stages (CS11) this labels almost exclusively neural crest cells in the developing CVG area, its’ expression in the OV starts to be detectable at CS12-13 (**Fig. 2D-F**).

### Induction of placode tissue from human iPSC

We then turned to a previously reported protocol to guide the differentiation of human iPSC towards otic fate (Koehler et al., 2017; Nie and Hashino, 2020) with the goal to comparing otic tissue derived from iPSC to the human samples. As in previous studies, different requirements for exogenous BMP4 have been reported to guide placode differentiation in hESC and hiPSCs, we tested varying concentrations of BMP4 (from 0.5ng/ml to 10ng/ml) (**Fig. 3 and Fig. S2**). Substantial morphological differences can be observed already at day 4 *in vitro* after exposure to BMP4 (**Fig. 3A**). The co-expression of the placodal marker SIX1 and Cadherin-1 (CDH1) was used to evaluate placode induction efficiency in a first set of experiments (**Fig. 3B-C and Fig. S2A**). BMP4 levels between 1 ng/ml and 2.5 ng/ml are required to generate placodal ectoderm (PE) with these lines. At optimal concentrations, 100% of the organoids display a SIX1+/CDH1+ placode-like layer on the surface of the aggregates (**Fig. 3C**). To confirm proper exposure to BMP4, samples were stained with an antibody recognizing the downstream target phospho-SMAD. Increasing amounts of recombinant BMP4 induce a dose-dependent phosphorylation of SMAD1-5-9 proteins (p-SMAD) (**Fig. 3D-E**).

**Figure 3:**
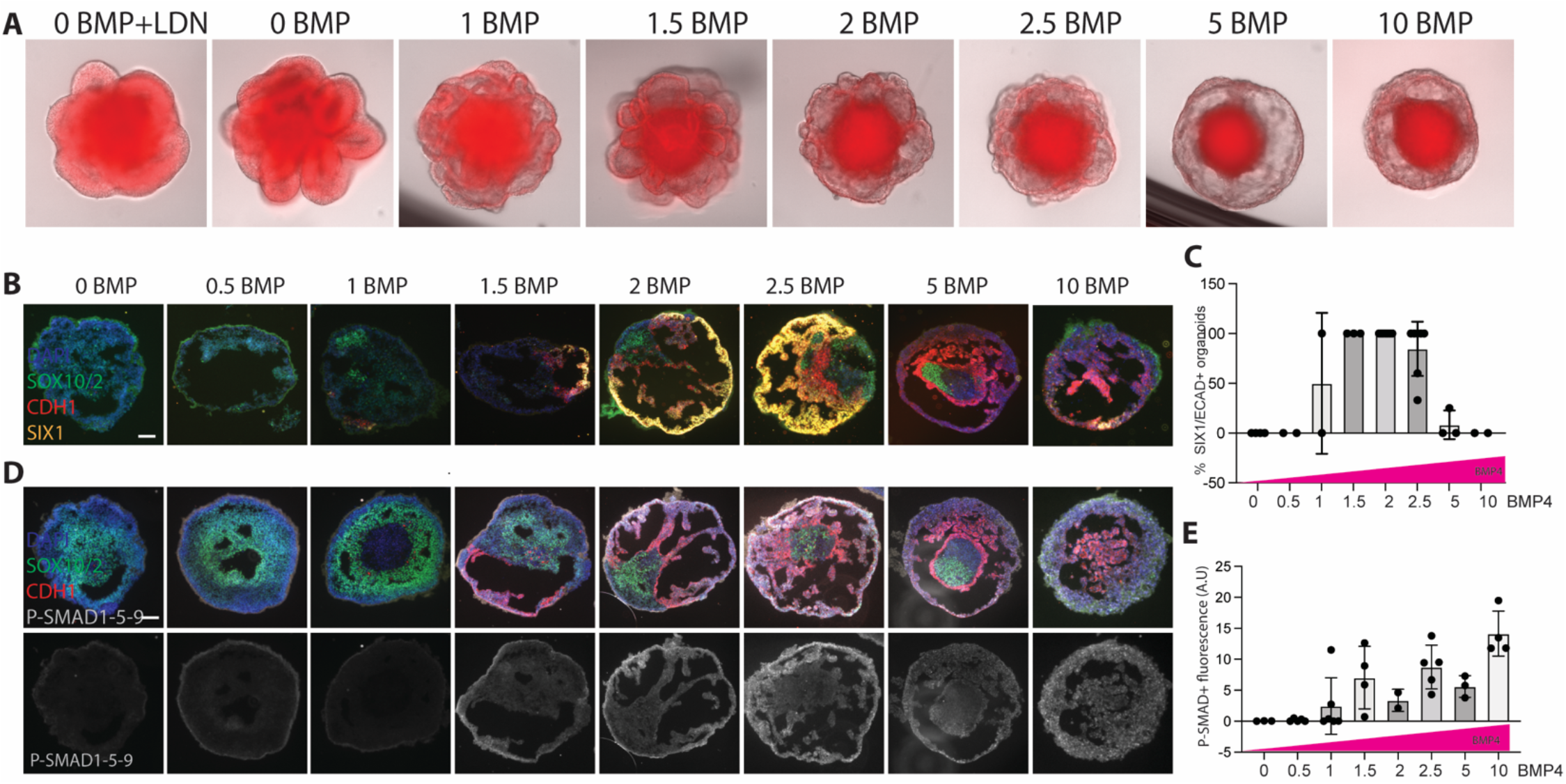
Placode induction is dependent on BMP concentration. **A-** iPSC aggregates derived from the LMNB-RFP iPSC line exposed to discrete BMP concentration from day 0 to day 3 ½ of culture imaged at day 4. **B-** iPSC aggregates derived from the SOX2-GFP iPSC line exposed to discrete BMP concentration from day 0 to day 3 ½ of culture. Organoids fixed at day 8 and immunostained for the placodal marker SIX1 and co-stained with CDH1 and SOX10. Scale bar 100μm. **C-** Quantification of the percentage of organoids with SIX1+CDH1+ domains at each BMP concentration. Average of 2-7 independent experiment with 2 cell lines (SOX2-GFP; PAX2-GFP iPSC). **D-** Same organoids as shown in B labelled with a phospho-specific antibody against P-SMAD1,5,9; CDH1 and SOX10. **E-** Image-based quantification of p-SMAD positive signal per organoids for all conditions. Average of 2 independent experiments with 2 cell lines (SOX2-GFP; PAX2-GFP iPSC).

To better characterize this system, we further analyzed the fate of the organoids cultured in the absence of exogenous BMP: BMP4 (0 ng/ml). Neither of the cell lines tested upregulates NNE/PE markers (AP2, CDH1, SIX1), instead we observed neuronal progenitors (SOX2, NEUROD1, Ki67), neurons (Cadherin-2 (CDH2), TUBB3) and glia (CDH2, SOX10) in untreated organoids (**Fig. S2C** see also data in **Fig. S7A-B** discussed later). In response to high levels of BMP (5-10ng/ml), the expression of GATA3, p63, and CDH1, is observed, suggesting that the NNE has committed to surface ectoderm/epidermal fate (**Fig. S2D**).

### Transcriptional characterization of the placodal tissue in IEOs

In order to characterize in depth the placodal tissue, as well as the cellular composition of the organoids at this stage, we employed single-cell RNA sequencing (scRNAseq). First, we assessed the percentage of epithelial cells differentiated at day 8 by flow cytometry analysis. As 40.57% +/- 9.4 (n=4 exp) of cells expressed the epithelial marker EPCAM, we decided to analyze whole organoids, expecting we would get enough coverage of the placode tissue of interest, as well as of the co-developed cell types. After data filtering (see method section) 15116 cells were analyzed from 200 pooled day 8 organoids derived from the SOX2-GFP iPSC line.

The two major cell populations that can be identified consist of an epithelial component (*EPCAM* and *CDH1* positive), representing *ca*. 70% of all cells, and a remaining mesenchymal/neuronal component (*EPCAM* and *CDH1* negative) (**Fig. 4A-E and Fig. S3)**. The two major groups of epithelial cell clusters include: placodal ectoderm-like (PE) cells and surface ectoderm-like cells (SE) (**Fig. 4A-B**). The PE cluster expresses the epithelial markers *EPCAM* and *CDH1*, in addition to *CDH2* and *SIX1* (**Fig. 4D-E and Fig. S3A**). Overall, this “pan-placodal” population represents 47% of all cells. Among the markers distinctive for PE cells identified in the scRNAseq experiment, we found *SPARCL1* and *BMP4* (**Fig. 4F**) Six PE subclusters can be distinguished by unsupervised clustering. Cells expressing posterior/otic-epibranchial placode (OEPD) markers (subclusters PE5, PE7, PE9) represent 21% of total cells (and 43% of the PE population) and express *PAX8.* In addition, these cells express *FGF3* and *FGF8.* Both genes were previously reported as important for otic development in different vertebrates (Hidalgo-Sanchez et al., 2000; Leger and Brand, 2002; Maroon et al., 2002; Zelarayan et al., 2007). We did not detect *PAX2* nor *SOX10* expression at day 8 in this cell population (**Fig. 4D-E**). Both genes become upregulated at later stages in the otic epithelium (**See Figure 5**). The clusters PE0 and PE1, include cells expressing OTX1 (12% of PE), OTX2 (16% of PE) and PAX3 (5% of PE), potentially representing anterior-placode (AP) cells at this stage. (Baker et al., 1999; Steventon et al., 2012) (**Fig. 4F**).

**Figure 4:**
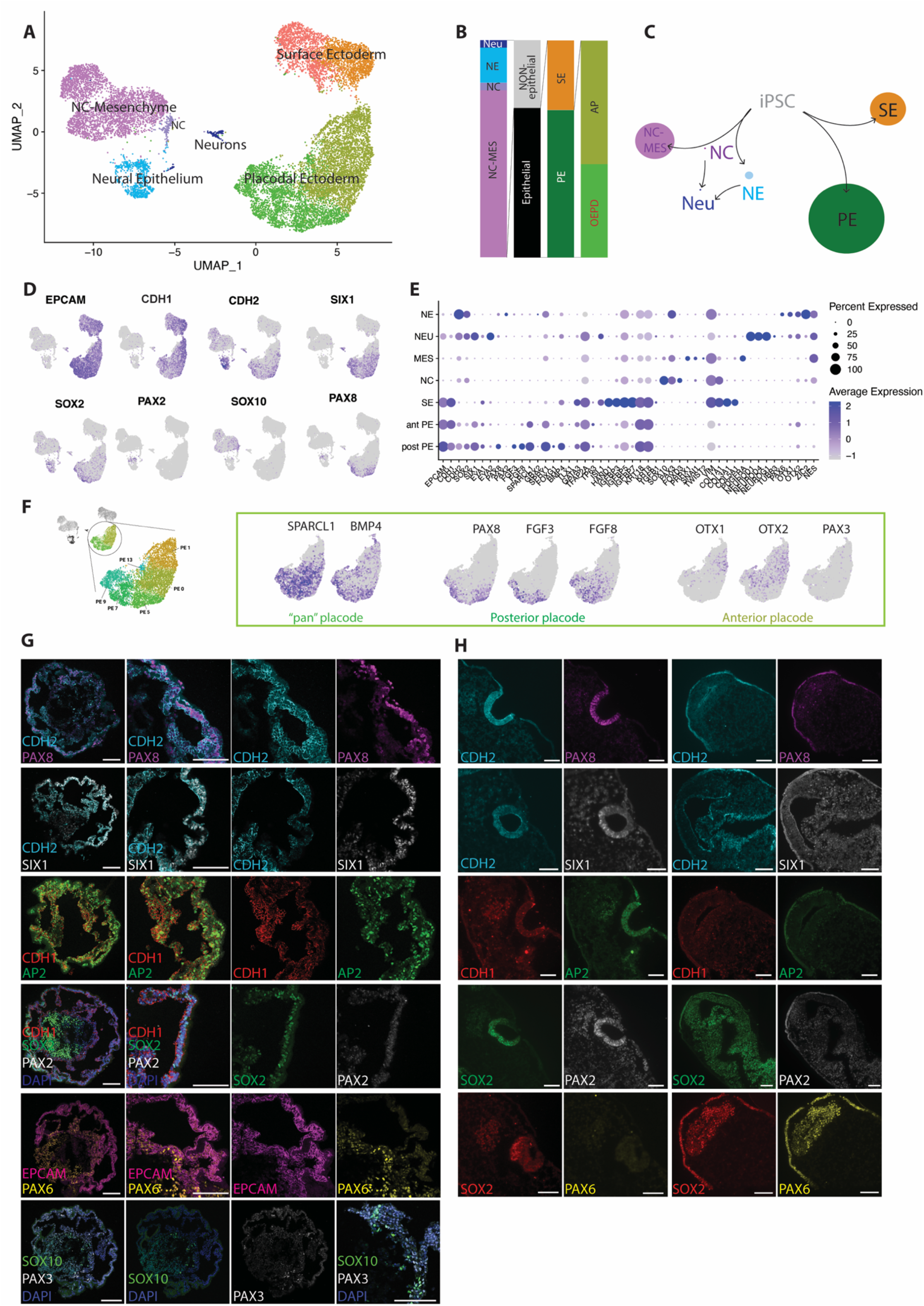
Comparison placodal tissue differentiated in IEO and CS11 samples. **A-** UMAP plot of day 8 scRNAseq data showing the identified cell clusters. Placodal ectoderm (PE; Green:); Surface ectoderm (SE; Orange); NC-Mesenchyme (NC-MES; Magenta); Neural Epithelium (NE; Turquoise:); Neural Crest (NC; Violet); Neurons (NEU; Blue:). **B-** Histogram plot showing the relative abundance of the different populations at day 8. OEPD: otic-epibranchial placode; AP: anterior placode. **C-** Schematic representation of the potential differentiation dynamics. Size of the circles proportional to the percentage of cells in the cluster, relative to total number of cells. **D**- UMAP plots with selected marker genes defining the PE cluster. **E-** Dot plot for selected markers for the main clusters. Abbreviations as above ; ant PE: anterior placode (PE0, PE1; PE13); post PE: posterior placode (PE5, PE7, PE9). **F-** UMAP plots for the PE population only (manual selection) 6 subclusters PE0; PE1; PE9; PE7; PE5; PE13 are identified by unsupervided clustering. ”pan placodal”, “anterior”, or “posterior” placode markers are shown. **G-** Organoid characterization at day 8. Scale bar= 100μm. Immunofluorescent staining characterization on consecutive sections was used to verify the expression/absence of the following markers: CDH1, CDH2, SIX1, AP2, PAX8, PAX2, SOX2, PAX6, SOX10, PAX3. **H-** Otocyst (left) and developing brain (right, same section) stage CS11 (E1026) immunostained for PAX8, SIX1, CDH1, AP2, SOX2 and PAX6. Scale bar= 100μm.

**Figure 5:**
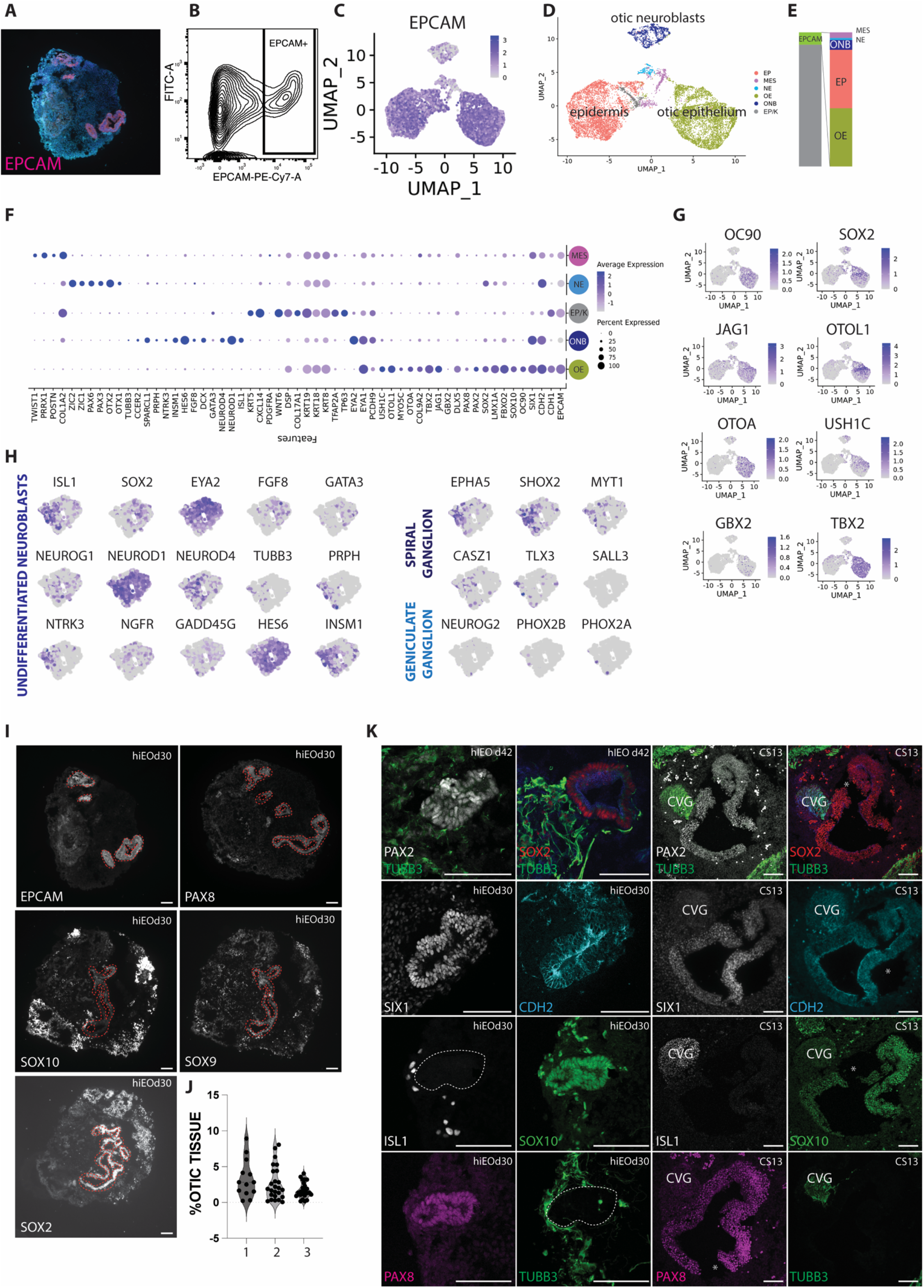
Comparison otic tissue differentiated in IEO and CS13 samples. **A-** EPCAM (magenta) and F-Actin (cyan) staining on an organoid section at day 30 of *in vitro* differentiation. **B-** Flow cytometry analysis of cells dissociated at day 26 of differentiation and stained for EPCAM-PeCy7. EPCAM+ cells (9.2% of the total) were sorted and processed for scRNAseq. **C**-UMAP plot of day 26 IEO scRNAseq showing expression of EPCAM in all cells analyzed. **D-** UMAP plot of day 26 IEO scRNAseq with cluster labeling. **E-** Percentage of the different cell populations identified by scRNAseq. **F-** Dot plot of the selected populations: Mesenchyme (MES, Magenta); Neural epithelium (NE, Turquoise); Epidermis/keratinocytes (EP/K, Gray); Otic neuroblasts (ONB, Blue); Otic epithelium (OE, Green). **G-** UMAP plots of selected genes expressed in the otic epithelium cluster. **H-** UMAP plots of selected genes labeling the otic neuroblasts cluster (manual selection). **I-** Characterization of otic vesicle in SOX2-GFP iPSC line at day 32 of differentiation. The otic markers EPCAM; PAX8; SOX10; SOX9 and SOX2 are shown. Scale bar 100μm. Vesicle areas is highlighted with a red contour. **J-** Quantification of the organoid area positive for each marker (n=3-4 sections of 3 organoid per experiment; 3 independent experiments). **K-** Otic vesicle at day 30-42 of differentiation (hIEOd30, hIEOd42) in comparison with CS13 embryo. Scale bar 100μm. *\* potential cutting artifact or tissue damage*.

The other cellular components detected in the data set include surface ectoderm-like cells (SE), 22% of the total cell number (**Fig. 4A-B**), expressing several NNE/epidermal markers: *CDH1, EPCAM, GATA3, TFAP2A, ISL1, HAND1, KRT18* and *KRT8* (**Fig. S3C**). Among the non-epithelial clusters, the following populations were found (shown in **Fig. S3C**): **i)** a neural crest (NC) population (1%), which we classified based on the expression of genes such as *SOX10, PAX3, PLP1 and FOXD3*. **ii) ;** NC-derived mesenchyme (Soldatov et al., 2019) (24%), which in addition to NC markers, shows upregulation of epithelial to mesenchymal transition (EMT) markers (*PRRX1, SNAI1, TWIST1/2* and *VIM*) (clusters MES4-MES6-MES8) **iii):** a small population of neurons (1%) positive for *NEUROD1, NEUROD4, NEUROG1, ISL1and TUBB3*, also expressing a number of placodal genes, suggesting they may be placode-derived new-born neurons (Durruthy-Durruthy et al., 2014). Finally, we also detected a cluster of *CDH2* and *SOX2* positive neural-epithelial cells (**iiii**) (5% of total), with two distinct gene expression sub-clusters, one with high expression of *PAX6 and OTX1/2* and the other expressing *PAX2* and *PAX8* (**Fig. S3C**).

Overall, the protocol yields a distinct cell population of placodal ectoderm cells and includes cells with a posterior/OEPD signature. The other populations derived in our cultures agree with the expected “off-target” tissue generated by the protocol (Koehler et al., 2017), possibly receiving insufficient or excessive BMP signal activation, as reported in **Fig. S2**.

### Placode characterization and comparison with human samples

To confirm the transcriptional data, we implemented immunostaining of consecutive sections and exploited tile-scanning to evaluate marker expression across conditions and time points (**Fig. S2B**). Already at day 4 or differentiation, the outer epithelial layer expresses CDH1, CDH2, TFAP2A (AP2) and SIX1 and lacks PAX2, PAX6 and SOX10 expression. Sparse SOX10+ NC cells can be detected in the core of the organoid (**Fig. S4A**). Using the SOX2-GFP reporter line, we observed that the core also expresses GFP. This could represent remaining SOX2+ pluripotent cells or neuronal progenitors. SIX1 expression is further maintained at days 8 and 12 on the surface of the aggregate (**Fig. S4B**). Starting at day 8, the otic marker PAX8 is upregulated in the outer layer of the organoids in cells that still co-express CDH1/CDH2/EPCAM/TFAP2A and SIX1(**Fig. 4G**). We could not convincingly detect PAX2 expression in the PE layer at this time point. PAX2 expression in the otic placode is also slightly delayed compared to PAX8 expression in mouse otocyst development (Bouchard et al., 2010). Some PAX6+ neural epithelial cells can be identified in the core of the organoids together with SOX10+ NC cells and SOX2+ neuronal progenitor cells (**Fig. 4G**). This pattern of marker expression is maintained up to day 12 (data not shown). We also evaluated the expression of SPARCL1, which indeed can be observed in the outer epithelial layer in day 8-12 organoids (**Fig. S4C**). Overall, the cellular composition identified by immunostaining matches the scRNAseq characterization, with the identified placodal layer positioned at the organoid surface.

We then stained CS11 embryos (days 24-25) for the same marker panel (**Fig. 4H).** The *in vivo* time point represents a slightly later developmental stage, nevertheless, many of the placodal markers were similarly expressed, including CDH1, CDH2, SIX1, PAX8 and AP2. PAX6 can be only detected in the developing brain and not in the otocyst (**Fig. 4H and S1B**). As shown in **Fig 2**, SOX10 staining at this stage mostly labelled NC cells in the region of the developing CVG, but not the epithelial components of the otocyst, as found in the organoids.

### Otic vesicle characterization

As a next step, we characterized by transcriptional profiling otic tissue differentiated within the organoid after 20-30 days of culture. Again, we first estimated the percentage of epithelial cells by staining IEO sections or dissociated cultures with an antibody recognizing EPCAM. Epithelial cells represented 8.6 % +/- 0.7 (n=2 experiments; day 27 and day 30) of total cells, when assessed by flow cytometry, and 5.7+/- 2.5 (n=2 experiments; day 30) by image analysis. To increase the yield of otic progenitors for single-cell analysis, we therefore decided to sort EPCAM-positive cells. We pooled together 140 organoids at day 26 of differentiation for dissociation to single cells. EPCAM-positive cells were sorted (**Fig. 5B**) and loaded on the 10X Chromium. Eventually 11793 cells were analyzed (**Fig. 5C-H and Fig. S5**). The great majority of cells express indeed EPCAM (**Fig. 5C**). Two major epithelial clusters can be identified in the dataset: **i**) Otic epithelium (OE) which represents 43.12% of all cells, and **ii)** epithelial cells with an epidermal-like signature (EP), representing 43.7% of the total (**Fig. 5D-E**). The otic epithelia derived *in vitro* co-express *CDH1* and *CDH2*, while the latter is absent in the EP clusters (**Fig. 5F and Fig. S5A**). OE further expresses high levels of *OC90, FBXO2, LMX1A,* and *TBX2*, as previously identified in mouse studies (Durruthy-Durruthy et al., 2014; Hartman et al., 2015; Kaiser et al., 2021; Sun et al., 2022) (**Fig. 5F-G and Fig. S5A, D**). The placodal genes *SIX1, DLX5* and *PAX8* are still expressed; *PAX2* and *SOX10* become now clearly upregulated (**Fig. 5F and Fig. S5D**). Interestingly, the expression of *SOX2* is limited to a subset of OE cells (59%), potentially indicating that both sensory and non-sensory domains are derived *in vitro*. The population of cells expressing high levels of *SOX2* also expresses *JAG1*. In addition, inner ear genes, such as *OTOL1, OTOA* and *USH1C* can also be identified in this cell population (**Fig. 5G).** Unsupervised clustering identified five distinct OE subclusters (**Fig. S5B**), we could however not differentiate them based on the expression of the above-mentioned marker genes.

Interestingly, we were able to distinguish a population of putative otic neuroblasts (ONB) (7.6% of total cell sequenced), expressing low levels of EPCAM, which could be co-isolated with this sorting approach (**Fig. 5F, H and Fig. S5A, D**). These cells express markers such as *ISL1, SOX2, EYA2, FGF8, GATA3, NEUROG1, NEUROD1* and *NEUROD4,* (Durruthy-Durruthy et al., 2014; Kaiser et al., 2021; Sun et al., 2022). In addition, ONBs start to express neuronal filaments like beta 3 tubulin (*TUBB3*) and peripherin (*PRPH*), and the neurotrophins receptors *NTRK3 and NGFR*. *GADD45G, HES6* and *INSM1*, recently identified by scRNAseq in the developing CVG in mice (Sun et al., 2022) are also expressed in the ONB population. Interestingly, markers identified in this latter study to distinguish spiral ganglion neurons (SGNs) from vestibular ganglion neurons are also expressed (*EPHA5; SHOX2; MYT1; CASZ1*) suggesting that the ONB population may resemble developing SGNs.

Finally, a small fraction of EPCAM negative cells was also detected, including neural epithelial cells (1.5% of total) and mesenchyme (4% of total) (**Fig. S5 A-C**).

As sorted cells processed for scRNAseq constituted in this experiment 9.2% of the total cell pool (**Fig. 5B**), we estimate OE cells represent on average 4% of the IEO composition at day 26.

To confirm these data, protein expression for OE markers was assessed by immunostaining and quantified across the three cell lines. We immunostained consecutive sections of organoids at day 30 for FBXO2, SOX10, SOX9, PAX2, PAX8, DLX5 and the epithelial markers EPCAM and CDH1 (**Fig. 5I Fig. S6** and data not shown). Image-based quantification at day 30 of differentiation shows that also in this case the otic tissue represents on average 2.4+/-2% of the whole organoid volume. Similar differentiation efficiency and marker expression were observed in the 3 different cell lines used in this study and different experiments (**Fig. 5J**).

Finally, we compared a subset of these markers between day 20-40 IEO and CS13 (day 28-32) samples (**Fig. 5K,** see also **Fig. 2).** PAX2 and SOX2 become now distinct in the IEO vesicle matching the *in vivo* expression. SIX1 and CDH2 are expressed both in CS13 samples as well as in d30 vesicles, confirming also the scRNAseq data. SOX10 at this time point becomes expressed in the OVs, besides the surrounding glia, while ISL1 is absent both in the IEO as well as in CS13 samples at the level of the otic epithelium. *In vivo* ISL1 is expressed in neuroblasts in the CVG and other cranial ganglia (glosso-pharyngeal ganglion and vagus nerve are shown in **Fig. S7D**). The population of ISL1 positive cells at CS13 expresses the highest level of TUBB3, indicating ISL1 positive cells are new-born neurons. This matches our transcriptional analysis, as well as the histology of IEO (**Fig. 5K**). At this time point *in vitro* otic vesicles are surrounded by neurons co-developed in culture. At earlier time points (d18), these are found organized in ganglia-like structures expressing ISL1 and being intermingled to SOX10+ and CD271+ cells (**Fig. S7 A-C**).

### Sensory vesicle maturation and comparison with the human inner ear

Starting from day 50-60 of IEO culture, hair cells develop within the organoids. These are organized in sensory epithelia intercalated to SOX2 positive supporting cells and receive innervation by co-developed neurons (Koehler et al., 2017; Nie and Hashino, 2020; Nie et al., 2022). Similar epithelial sensory patches have been obtained with the 3 cell lines tested in this study (**Fig. 6A-C and Fig. 7A**). We collected organoids at day 60 from two independent experiments for scRNAseq (see method section). In both experiments EPCAM positive cells were sorted (**Fig. S8A**). We sequenced 7054 cells in experiment 1 and 6440 cells experiment 2. The two data sets were then integrated for further analysis (Stuart et al., 2019). Despite sorting epithelial cells, some EPCAM negative cells, including mesenchyme, neural epithelium and neurons have also been identified, likely due to an non-stringent gating strategy in experiment 1 (**Fig S8D**). These populations were not further analyzed.

**Figure 6:**
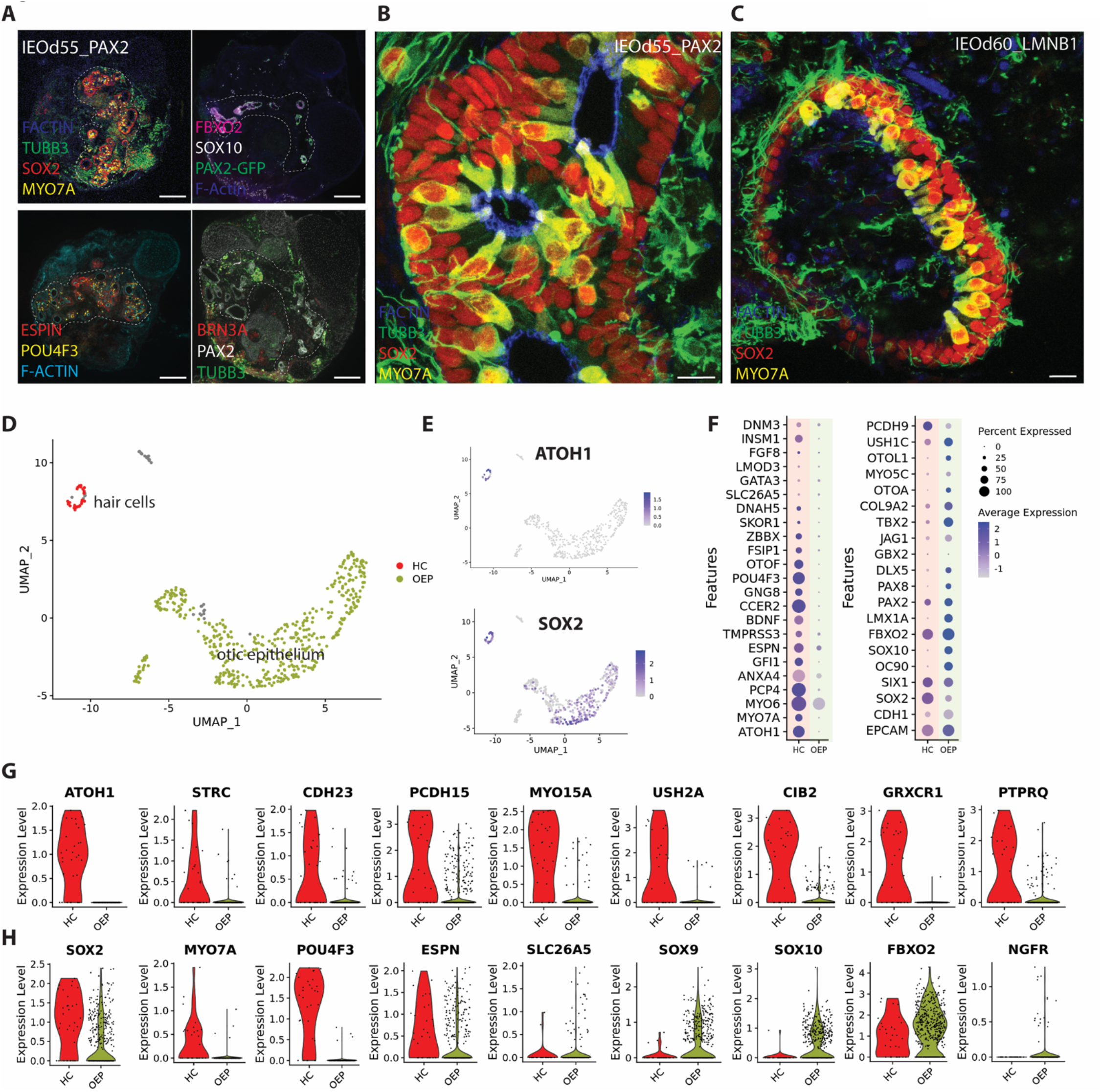
Characterization HC and sensory epithelial in IEOs at day 60. **A-** Characterization of IEO at day 55 of differentiation (PAX2-GFP iPSC line). Different sections of the same organoids are shown. Otic marker positive areas contoured with white-dashed line. Scale bars 100μm. **B-** Sensory vesicle derived from the PAX2-GFP iPSC line (IEO d55) and **C-** LMNB1-RFP iPSC line (IEOd60). Hair cell (MYO7A+/SOX2+); Supporting cells (SOX2+/MYO7A-) neurons (TUBB3) and F-Actin staining. Scale bars 10μm. **D-** UMAP plot of selected OEP (green) and HC (red) clusters. **E-** UMAP plots of the selected cells showing *ATOH1* and *SOX2* expression **F-** Dot plot with relative expression of known marker genes in the selected OEP and HC cluster. **G** and **H-** Violin plots showing expression levels of selected genes in HC and OEP.

**Figure 7:**
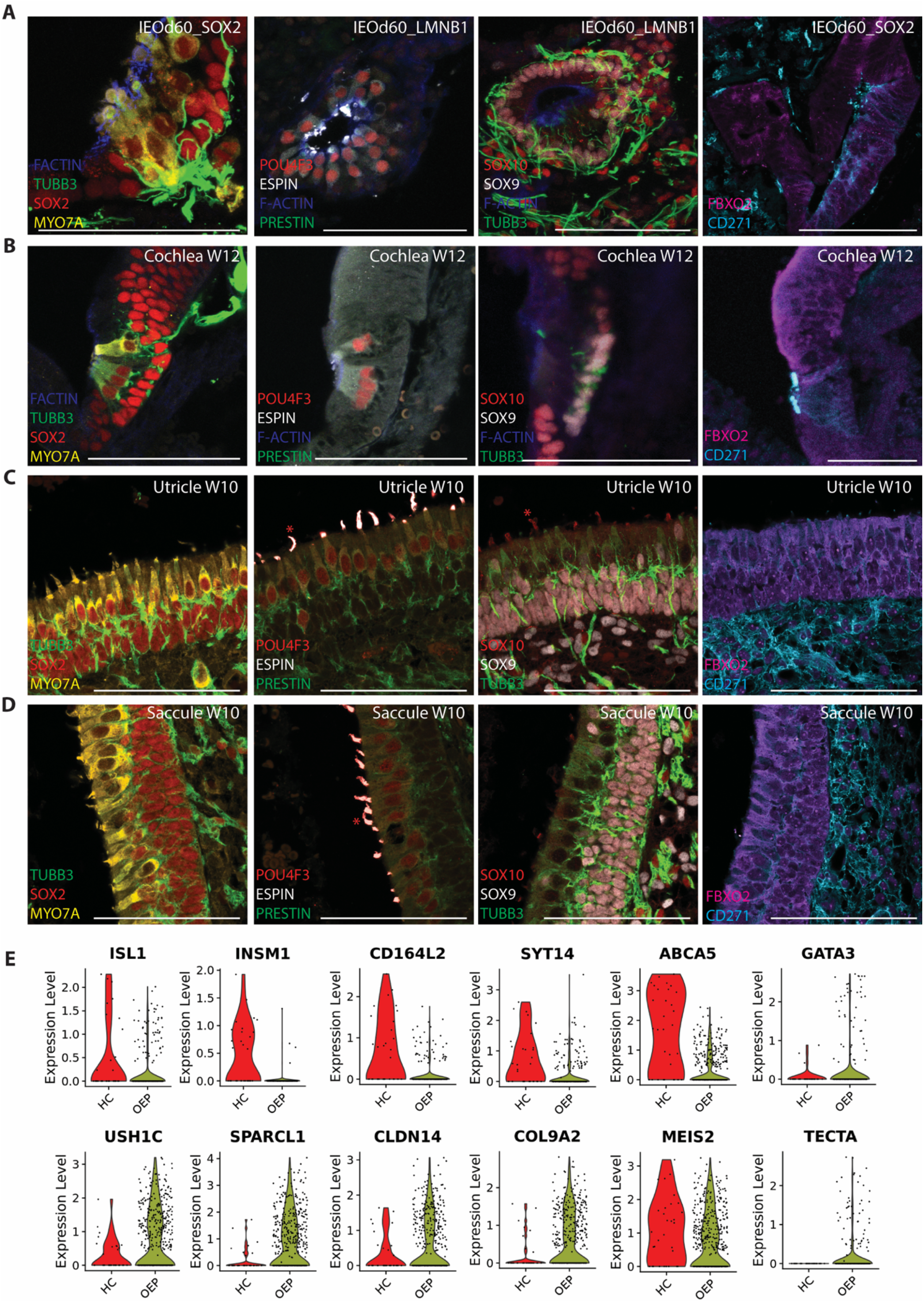
Comparison organoid marker expression and human fetal cochlea, utricle and saccule. **A-** Representative images of IEO sensory vesicles from two different cell lines (SOX2-GFP iPSC and LMNB1-RFP iPSC) at day 60 of differentiation. **B-** Cochlea sections at week 12 of development (sample E1291). **C-** Utricle sections at week 10 of development (sample EF1). **D-** Saccule sections at week 10 of development (sample EF1). Scale bar 100μm. *\* asterisks indicate cross-reactivity of the goat anti mouse IgG1 antibody (used for POU4F3 and SOX10 staining) with hair bundles. This has been detected in different staining, only on paraffin embedded sections.* **E-** Violin plots showing expression levels in HC and OEP for selected genes.

Among the epithelial components (EPCAM+), epidermal cells (EP) form the most abundant population. A distinct cluster of otic epithelial cells (OEP, 498 cells) can be identified, expressing very specifically *OC90, FBXO2, SOX10, OTOL1* and *USH1C* (**Fig. S8E**). As putative hair cells did not form a separate cluster, we selected cells using the module score feature of Seurat (see methods), using gene sets expressed by HC and OEP (**Table 2**) and selected for further analysis these two populations (**Fig. 6D-H**). The identified hair cells not only express newborn-HC markers (Burns et al., 2015; Kolla et al., 2020), including *ATOH1, POU4F3, GFI1, CCER2* and *GNG8,* but also genes associated to HC maturation such as *STRC, CDH23, PCDH15, ESPN, TMPRSS3 and OTOF* (**Fig. 6F-H and Fig. S8G, I**). Among the top differentially expressed genes in HC versus the OEP, we have identified *MYO15A, USH2A, CIB2, GRXCR1* and *PTPRQ*. Many of these genes have been associated to hearing loss before (www.hereditaryhearingloss.org) (**Fig. S8I**).

We then compared the *in vitro* generated sensory epithelia by transcriptional profiling (**Fig. 6H**) and histological assessment (**Fig. 7A**), to human specimens collected at W12 of development (cochlear samples, **Fig. 7B**) or at W10 (vestibular samples: utricle **Fig. 7C,** and saccule **Fig. 7D**). Comparable protein expression and protein localization are observed for MYO7A, ESPIN and POU4F3 in the three sensory epithelia analyzed, as well as in the IEOs (**Fig. 7A-D**). SOX2 is still expressed in hair cells at this stage (Locher et al., 2013; Roccio et al., 2018). We did not observe expression of Prestin in OHC in the cochlear tissue at W12. A similar faint immunoreactivity was detected in the IEO vesicles. In line with this, *SLC26A5*, encoding for Prestin, was expressed only by a few cells (**Fig. 6H**). Prestin expression has been reported in later developmental stages than what analyzed here, starting only at p9-10 in mice (Hang et al., 2016). Hair bundles, though clearly expressing Espin are immature at these stages, both *in vivo* as well as *in vitro* (**Fig. 7A-D**).

Supporting cells express beside SOX2, also SOX9 and SOX10 in IEOs and primary tissues. SOX10 positive cells can be detected also below the sensory epithelia in the vestibular organs and in IEOs (**Fig. 7 D-E**), likely labelling glial cells. FBXO2 staining can be observed in the cochlear duct, in the vestibular epithelia and IEOs, but the expression is not restricted to sensory areas, as shown in the murine inner ear (Hartman et al., 2018). We previously reported the expression of CD271 (NGFR), in the developing human cochlear prosensory domain (Roccio et al., 2018). Interestingly a similar pattern of NGFR expression could be observed in some of the vesicles in the IEOs (**Fig. 7B**).

Finally, *ISL1* can be identified both in hair cells and in supporting cells (**Fig. 7E**). By histological analysis of the IEOs we found day 60 sensory vesicles expressing ISL1. A similar expression pattern can be observed in the sensory epithelia of utricle, saccule and cochlea (**Fig. S9A**).

To assess if we could further distinguish different HC identities, we analyzed the expression of genes reported in studies comparing cochlear and vestibular sensory epithelia (Burns et al., 2015; Elkon et al., 2015; Scheffer et al., 2015; Wilkerson et al., 2021) (**Fig. 7E**). We identify *INSM1,* previously described in cochlear OHC (Elkon et al., 2015; Wiwatpanit et al., 2018) as well *CD164L2, SYT14* and *ABCA5,* recently described in adult human vestibular hair cells (Wang et al., 2022). The transcription factor *GATA3*, presumably associated to cochlear identity, is instead expressed at low levels in a small number of hair cells, but present in OEP cells, matching our histological characterization (**Fig. S9B**). Within the OEP subcluster, we detected expression of *USH1C, SPARCL1, CLDN14* and *COL9A2. MEIS2,* previously described in vestibular epithelia, and TECTA, identified in cochlear epithelia (Wilkerson et al., 2021), are also expressed. Overall, the maturity of the IEO-derived hair cells and supporting cells at day 60 of *in vitro* differentiation corresponds to W10-12 human fetal development. We were however not able to provide a clear distinction of vestibular or cochlear identity at this stage.

## Discussion

### 1. Benchmarking human iPSC derived inner ear organoids

The restricted access to human inner ear tissue is a primary limiting factor to study its development and to validate therapeutic strategies for hearing and balance restoration (Roccio, 2021). In this respect, stem cell-based models, such as the inner ear organoids described here, offer a promising alternative to this problem. In contrast to models which rely on human somatic cells (Chen et al., 2007; Roccio et al., 2018; Senn et al., 2019; Taylor et al., 2015) or trans-differentiation strategies (Costa et al., 2015; Menendez et al., 2020; Noda et al., 2018), the use of pluripotent cells as starting point enables to study early otic developmental stages. Furthermore, by generating complex sensory units, including sensory and non-sensory epithelia, neurons and surrounding otic mesenchyme, IEOs offer a platform to study more complex aspects of organ physiology and pathology as compared to monotypic cultures.

Here, we have evaluated in detail a previously reported protocol to guide iPSC through consecutive steps of otic development (Koehler et al., 2017; Nie and Hashino, 2020) and compared, for the first time, the *in vitro*-derived otic tissue to early human embryos. This represents a first step towards the benchmarking of IEOs. Similar studies have been performed with brain organoids (Camp et al., 2015; Pollen et al., 2019) retinal organoids(Cowan et al., 2020; Wahle et al.) pancreatic organoids (Goncalves et al., 2021) to name a few examples, with the complementary goal to create developmental human cell atlases (Haniffa et al., 2021). IEO are a relatively new addition to the catalogue of *in vitro* derived organ models. Our work, together with other recent unpublished studies, aim at providing a robust foundation for broader application of IEOs (Moore et al., 2022; Steinhart et al.; van der Valk et al., 2022).

While we provide a comparison using immunostaining of tissue sections, we have not been able to obtain and to process freshly isolated tissue from early embryos (CS11-13) for a comparative gene expression analysis using scRNAseq. This remains a limitation of this study and potentially a roadblock for similar endeavors. Not only given the difficulty in obtaining these early samples, but also due to the small number of cells forming the otic vesicle at this stage, making their identification in unsorted single cell embryo data sets very hard or impossible (Xu et al., 2022).

### 2. BMP signaling in placode development

The IEO protocol was developed with the goal to recapitulate otic development *in vitro* using consecutive addition of small molecules and growth factors (Koehler et al., 2013; Koehler et al., 2017). We have characterized three major stages of the induction protocol: placode differentiation, otic vesicle formation and finally, sensory epithelia maturation.

The first step is triggered in iPSC aggregates cultured in presence of Matrigel^TM^ and a TGFβ inhibitor, by exposure to low levels of FGF2 and BMP4. BMP is a potent inhibitor of neural fate and plays a crucial role for ectoderm differentiation towards non-neural fate (Wilson and Hemmati-Brivanlou, 1995). In the first 4 days of culture, BMP4 triggers the differentiation of NNE and PE on the surface of the iPSC aggregates. BMP signaling is subsequently inhibited, and high levels of FGF2 and WNT signaling are used to promote otic placode fate (Martin and Groves, 2006; Zelarayan et al., 2007). Optimization of the BMP4 level is a critical factor for successful placodal induction in IEOs in all the iPSC lines tested here. This requires fine tuning for each line, to account for endogenous morphogen production. In some lines (Koehler et al., 2017; Nie et al., 2022), or other studies using 2D approaches (Dincer et al., 2013; Ealy et al., 2016), no exogenous BMP was used to promote placodal fate. Experiments performed with hESC on micropatterns, point to co-integration of BMP and WNT signaling to induce placode differentiation (Britton et al., 2019). In presence of BMP, WNT signaling favors the co-development of NC from ectodermal progenitors. Inhibition of WNT with the inhibitor IWP2 on the other hand, enhanced placodal fate even with low BMP4 supplementation. The co-development of NC and placode in IEO suggests the presence of endogenously secreted WNTs, and possibly BMP. This agrees with the extensive NC differentiation observed in absence of added BMP4 (**Fig. S2).**

The localization of phosphorylated SMAD proteins observed in response to exogenous BMP4 (**Fig. 3D)** suggests a differential activation of BMP signaling across the lateral-medial axes of the organoids depending on the dose of the morphogen. High doses activate SMAD signaling also in the organoid core, while low levels (1.5-2.5 ng/ml), induce SMAD-phosphorylation only at the external NNE layer. This differential pattern of BMP activation may enable the co-differentiation of NC and NC-derived mesenchyme. Even though considered “off-target” tissues, these may provide the proper context for subsequent morphogenetic events leading to otic vesicle formation *in vitro*. We compared IEOs at the placode induction stage (d8-12) with human samples at CS11 (d24-25). Even though not perfectly matching in terms of developmental time points, most of the placodal markers were conserved and similarly expressed. PAX2, already expressed *in vivo* in the invaginating otic placode was instead not yet detected in organoids at day 8, but appeared in IEOs starting at day 20.

### 3. Otic epithelia and neuroblasts differentiation

The second stage we characterized spans between day 20-40 of *in vitro* differentiation, during which otic epithelia and neuroblasts arise. Both cell types were indeed identified in the day 26 RNAseq data set. IEO-derived otic epithelia and neuroblasts express markers previously found in the developing murine otocysts (Durruthy-Durruthy et al., 2014; Hartman et al., 2015), and based on our histological characterization match the CS13 stage of development.

Our characterization of IEO-derived otic epithelium corroborates recently reported findings (Nie et al., 2022). IEO-neuroblasts, besides general undifferentiated markers express genes such as *CASZ1, EPHA5 and SHOX2* and lack expression of *SAL3*, matching the pattern recently reported in the developing cochlear ganglion (Sun et al., 2022). Furthermore, they lack expression of geniculate ganglion neurons genes such as *PHOX2B, PHOX2A* and *NEUROG2*, suggesting these neuroblasts may resemble spiral ganglion neurons precursors.

In this study we did not assess if these ONB eventually developed in SGNs. Our analysis at later time points focused on the sensory epithelia components. The neuronal population identified at day 60 (**Fig. S8**), after EPCAM sort, may represent only a fraction of less differentiated cells. We have therefore not analyzed it in detail. Transcriptional profiling of whole organoids is starting to shed light on the neuronal populations derived in IEOs (Steinhart et al.). Nevertheless, in absence of a complete transcriptional atlas for cranial ganglia, DRGs and concomitant analysis of CNS-neurons, we feel it is currently not possible to assign identity to these populations unequivocally.

### 4. Hair cell maturation

*In vivo* differentiation and maturation of hair cells in the inner ear sensory patches is not synchronous and it happens over the course of a few weeks in humans. Hair cells in the vestibular organs develop prior to hair cells in the cochlea (*circa* 2 weeks). Cells at the base of the cochlea develop prior to cells in the apical turn, and inner hair cells differentiate prior to outer hair cells. The whole process spans between W8 and W13 (Locher et al., 2013; Roccio et al., 2018).

Our comparison between IEO-derived sensory vesicles at day 60 *in vitro* and the human inner ear, shows a similar marker expression as for the fetal cochlea at W12 and the human fetal utricle and saccule at W10. This is the case both for hair cells and supporting cell markers.

Our transcriptomic data shows that the hair cells derived by day 60 *in vitro* express “general” hair cell markers. At this time point, the gene signature defining sub-classes of hair cells (cochlear versus vestibular) is not yet clear. We found in fact genes previously associated with both types of sensory epithelia. This could indicate either that both types differentiate, but given the timing, a more substantial number of vestibular hair cells would be present at early stages. Alternatively, the cues provided for otic induction, in particular WNT activation, may be skewing the culture to dorsal/vestibular fate. Recent work exploiting scRNAseq to characterize hair cells from WT or CDH7-mutant iPSC lines, suggest that the majority of cells derived *in vitro* after 70 days has a vestibular signature (Nie et al., 2022). Additional activation of SHH signaling and inhibition of WNT during otic vesicle maturation seem to promote cochlear fate. This becomes however apparent only in long term cultures >140 days (Moore et al., 2022).

While the number of studies focused on the molecular characterization of inner ear sensory epithelia has increased with the advent of single cell genomic (Burns et al., 2015; Li et al., 2018; Nie et al., 2022; Petitpre et al., 2022; Ranum et al., 2019; Sun et al., 2022; Waldhaus et al., 2015; Wilkerson et al., 2021), a comprehensive and simultaneous comparison of the different sensory epithelia and neurons, across developmental stages is currently lacking. The specificity and selectivity of many genes used as “markers” to each organ, and in particular for human tissues, is not yet robustly established.

GATA3 for example, is initially broadly expressed in the otocyst and then acts as key regulator of SGN and cochlear prosensory domain maturation in the mouse (Luo et al., 2013). While GATA3 has been identified in many studies as enriched in the cochlea (Cai et al., 2015; Elkon et al., 2015; Scheffer et al., 2015; Wilkerson et al., 2021) it has been observed also in the vestibular epithelia in the human cochlea (Johnson Chacko et al., 2020). Here we found low levels of GATA3 expression in a few hair cells. High *GATA3* expression was found instead in a subset of the OEP cells. These co-express *GATA2, BMP4, GJB2*, *GJB6* and *ISL1*. While this cluster seem to match the gene expression of non-sensory cochlear floor regions (lateral domain), both a higher number of cells and better references for cochlear and vestibular epithelia would be needed to conclude about the true nature of these cell types.

### 5- IEO characterization by scRNAseq strength and limitations

The use of a high number of “unselected” and unsorted organoids for scRNAseq is in principle an ideal tool to obtain an unbiased representation of the global efficiency of the induction protocol (Steinhart et al.; van der Valk et al., 2022). Nevertheless, less abundant cell types can be missed with this approach. Sorting strategies may allow to circumvent this issue.

At the placodal stage (day 8 of differentiation) posterior placodal cells represent in this study 21% of total cells. PE is localized at the external layer of the aggregates. It is therefore possible that the relative percentage of these cells may be limited by the aggregate size. In a subsequent step we have identified 4-5% of cells differentiating to otic epithelium and an additional 0.7% of cells representing otic neuroblasts. At this step, we relied on EPCAM expression and FACS sorting to enrich for otic progenitors. However, we did not collect the negative fractions, limiting our assessment of the full cellular composition of the organoids at this stage.

Our analysis at later time points (day 60), resulted in the identification of a small population of hair cells; however, by our estimation, the number of hair cells in our transcriptomic data set is inferior to what we observed in the histological analysis of the organoids. We suspect this may be due to problems triturating organoids to single cells, especially the compact epithelia we observe at day 60. Cell preparation methods are known to introduce biases in the abundancy of different cellular populations (Denisenko et al., 2020; Uniken Venema et al., 2022). In the absence of fluorescent reporters marking hair cells, the isolation of this specific population is hard to optimize. Fluorescent reporter lines for hair cells, supporting cells or neuroblasts, or additional surface markers to purify and enrich for low-abundance cells, should provide better coverage of these cell types in future studies.

## Outlook and conclusions

This study provides the first unique molecular analysis of otocyst development in human embryos, serving as a reference for further developmental studies. We demonstrate that the current state-of-the-art IEO protocol generates cell types that match, in terms of marker expression, human otic progenitors and follow similar developmental timings as the *in vivo* counterpart. IEOs could, therefore, significantly contribute to informing human otic development. Alternative strategies, possibly deviating from the developmental trajectory, may allow for further enrichment of the cells of interest at the expense of other tissues (Dincer et al., 2013; Qi et al., 2017).

Some level of variability nevertheless remains in IEO generation and final sensory epithelia maturation. Optimizing culture conditions, for example, using bioreactors or automated systems for medium exchange may eventually enable standardization and further enhance reproducibility for generating functionally mature cell types. The current model consists of closed vesicles or tubules that may not experience the same degree of fluid-induced shear stress, nor vibration-induced maturation that the inner ear experiences. The lack of these mechanical cues could limit the proper maturation of the different cell types. One could speculate that more advanced culture systems, such as organs-on-a-chip, may provide models to dissect the contribution of different factors to cell maturation.

## Acknowledgements

We thank the staff of the Flow cytometry core facility of the University of Zurich (Dr. Mario Wickert, Dr. Tatiane Gorski and Dr. Philip Schätzle) for their professional help and flexibility with sorting experiments. We thank the staff of the Genomic core facility, in particular Dr, Daniel Ehrsam and Dr. Qin Zhang for their help with sample preparation. In addition, we thank Edward van Beelen, from the Otobiology Lab, LUMC, for fetal specimen collection. Further, we thank Prof. Dr. Christian Grimm from the department of Ophthalmology, USZ, for access to the cryostat and Dr. Melanie Generali, from the iPSC core facility of the University of Zurich and IREM institute for help with iPSC cultivation. Image acquisition was performed using microscopes of the imaging core facility of the University of Zurich. The human PAX2-GFP iPSC line was generated within a previous EU-FP7-funded project (www.otostem.org). We thank Prof. Dr. Hubert Löwenheim and all the consortium members for sharing the line.

## Authors contribution

Data generation (D.D; S.A.J; V.V, M.R. W.H.V, A.E) data analysis (D.D; S.A.J; V.V; M.R; H.R, J.M.C), data interpretation (D.D; S.A.J; J.Z., M.R. K.R.K) human sample collection (H.R.W, S.S; H.L; W.H V; M.R), manuscript writing (S.A.J; M.R). DD and S.A.J have contributed equally to the study.

## DATA Accessibility

The single cell sequencing data generated by this study and presented in the publication have been deposited in NCBI’s Gene Expression Omnibus (Edgar et al., 2002) and are accessible through GEO Series accession number GSE229148 (https://www.ncbi.nlm.nih.gov/geo/query/acc.cgi?acc= GSE229148). Codes available on request.

## REFERENCES

Adameyko, I., Lallemend, F., Furlan, A., Zinin, N., Aranda, S., Kitambi, S. S., Blanchart, A., Favaro, R., Nicolis, S., Lubke, M., et al. (2012). Sox2 and Mitf cross-regulatory interactions consolidate progenitor and melanocyte lineages in the cranial neural crest. Development 139, 397–410.

Alsina, B. and Whitfield, T. T. (2017). Sculpting the labyrinth: Morphogenesis of the developing inner ear. Semin Cell Dev Biol 65, 47–59.

Appler, J. M. and Goodrich, L. V. (2011). Connecting the ear to the brain: Molecular mechanisms of auditory circuit assembly. Progress in neurobiology 93, 488–508.

Baker, C. V., Stark, M. R., Marcelle, C. and Bronner-Fraser, M. (1999). Competence, specification and induction of Pax-3 in the trigeminal placode. Development 126, 147–156.

Bok, J., Dolson, D. K., Hill, P., Ruther, U., Epstein, D. J. and Wu, D. K. (2007). Opposing gradients of Gli repressor and activators mediate Shh signaling along the dorsoventral axis of the inner ear. Development 134, 1713–1722.

Bouchard, M., de Caprona, D., Busslinger, M., Xu, P. and Fritzsch, B. (2010). Pax2 and Pax8 cooperate in mouse inner ear morphogenesis and innervation. BMC Dev Biol 10, 89.

Britton, G., Heemskerk, I., Hodge, R., Qutub, A. A. and Warmflash, A. (2019). A novel self-organizing embryonic stem cell system reveals signaling logic underlying the patterning of human ectoderm. Development 146.

Burns, J. C., Kelly, M. C., Hoa, M., Morell, R. J. and Kelley, M. W. (2015). Single-cell RNA-Seq resolves cellular complexity in sensory organs from the neonatal inner ear. Nat Commun 6, 8557.

Cable, J., Barkway, C. and Steel, K. P. (1992). Characteristics of stria vascularis melanocytes of viable dominant spotting (Wv/Wv) mouse mutants. Hear Res 64, 6–20.

Cai, T., Jen, H. I., Kang, H., Klisch, T. J., Zoghbi, H. Y. and Groves, A. K. (2015). Characterization of the transcriptome of nascent hair cells and identification of direct targets of the Atoh1 transcription factor. J Neurosci 35, 5870–5883.

Camp, J. G., Badsha, F., Florio, M., Kanton, S., Gerber, T., Wilsch-Brauninger, M., Lewitus, E., Sykes, A., Hevers, W., Lancaster, M., et al. (2015). Human cerebral organoids recapitulate gene expression programs of fetal neocortex development. Proc Natl Acad Sci U S A 112, 15672–15677.

Chen, W., Cacciabue-Rivolta, D. I., Moore, H. D. and Rivolta, M. N. (2007). The human fetal cochlea can be a source for auditory progenitors/stem cells isolation. Hear Res 233, 23–29.

Costa, A., Sanchez-Guardado, L., Juniat, S., Gale, J. E., Daudet, N. and Henrique, D. (2015). Generation of sensory hair cells by genetic programming with a combination of transcription factors. Development 142, 1948–1959.

Cowan, C. S., Renner, M., De Gennaro, M., Gross-Scherf, B., Goldblum, D., Hou, Y., Munz, M., Rodrigues, T. M., Krol, J., Szikra, T., et al. (2020). Cell Types of the Human Retina and Its Organoids at Single-Cell Resolution. Cell 182, 1623–1640 e1634.

de Bakker, B. S., de Jong, K. H., Hagoort, J., de Bree, K., Besselink, C. T., de Kanter, F. E., Veldhuis, T., Bais, B., Schildmeijer, R., Ruijter, J. M., et al. (2016). An interactive three-dimensional digital atlas and quantitative database of human development. Science 354.

Denisenko, E., Guo, B. B., Jones, M., Hou, R., de Kock, L., Lassmann, T., Poppe, D., Clement, O., Simmons, R. K., Lister, R., et al. (2020). Systematic assessment of tissue dissociation and storage biases in single-cell and single-nucleus RNA-seq workflows. Genome Biol 21, 130.

Dincer, Z., Piao, J., Niu, L., Ganat, Y., Kriks, S., Zimmer, B., Shi, S. H., Tabar, V. and Studer, L. (2013). Specification of functional cranial placode derivatives from human pluripotent stem cells. Cell Rep 5, 1387–1402.

Durruthy-Durruthy, R., Gottlieb, A., Hartman, B. H., Waldhaus, J., Laske, R. D., Altman, R. and Heller, S. (2014). Reconstruction of the mouse otocyst and early neuroblast lineage at single-cell resolution. Cell 157, 964–978.

Ealy, M., Ellwanger, D. C., Kosaric, N., Stapper, A. P. and Heller, S. (2016). Single-cell analysis delineates a trajectory toward the human early otic lineage. P Natl Acad Sci USA 113, 8508–8513.

Edgar, R., Domrachev, M. and Lash, A. E. (2002). Gene Expression Omnibus: NCBI gene expression and hybridization array data repository. Nucleic Acids Res 30, 207–210.

Elkon, R., Milon, B., Morrison, L., Shah, M., Vijayakumar, S., Racherla, M., Leitch, C. C., Silipino, L., Hadi, S., Weiss-Gayet, M., et al. (2015). RFX transcription factors are essential for hearing in mice. Nat Commun 6, 8549.

Evsen, L., Sugahara, S., Uchikawa, M., Kondoh, H. and Wu, D. K. (2013). Progression of neurogenesis in the inner ear requires inhibition of Sox2 transcription by neurogenin1 and neurod1. J Neurosci 33, 3879–3890.

Evtouchenko L, S. L., Spenger C, Dreher E, Seiler RW. (1996). A mathematical model for the estimation of human embryonic and fetal age. Cell Transplantation 4, 453–464.

Germain, P., Lun, A., Macnair, W. and Robinson, M. (2021). Doublet identification in single-cell sequencing data using scDblFinder. https://f1000research.com/articles/10-979/v1.

Goncalves, C. A., Larsen, M., Jung, S., Stratmann, J., Nakamura, A., Leuschner, M., Hersemann, L., Keshara, R., Perlman, S., Lundvall, L., et al. (2021). A 3D system to model human pancreas development and its reference single-cell transcriptome atlas identify signaling pathways required for progenitor expansion. Nat Commun 12, 3144.

Groves, A. K. and Fekete, D. M. (2012). Shaping sound in space: the regulation of inner ear patterning. Development 139, 245–257.

Gu, R., Brown, R. M., 2nd, Hsu, C. W., Cai, T., Crowder, A. L., Piazza, V. G., Vadakkan, T. J., Dickinson, M. E. and Groves, A. K. (2016). Lineage tracing of Sox2-expressing progenitor cells in the mouse inner ear reveals a broad contribution to non-sensory tissues and insights into the origin of the organ of Corti. Dev Biol 414, 72–84.

Hang, J., Pan, W., Chang, A., Li, S., Li, C., Fu, M. and Tang, J. (2016). Synchronized Progression of Prestin Expression and Auditory Brainstem Response during Postnatal Development in Rats. Neural Plast 2016, 4545826.

Haniffa, M., Taylor, D., Linnarsson, S., Aronow, B. J., Bader, G. D., Barker, R. A., Camara, P. G., Camp, J. G., Chedotal, A., Copp, A., et al. (2021). A roadmap for the Human Developmental Cell Atlas. Nature 597, 196–205.

Hao, Y., Hao, S., Andersen-Nissen, E., Mauck, W. M., 3rd, Zheng, S., Butler, A., Lee, M. J., Wilk, A. J., Darby, C., Zager, M., et al. (2021). Integrated analysis of multimodal single-cell data. Cell 184, 3573–3587 e3529.

Hartman, B. H., Bscke, R., Ellwanger, D. C., Keymeulen, S., Scheibinger, M. and Heller, S. (2018). Fbxo2(VHC) mouse and embryonic stem cell reporter lines delineate in vitro-generated inner ear sensory epithelia cells and enable otic lineage selection and Cre-recombination. Dev Biol 443, 64–77.

Hartman, B. H., Durruthy-Durruthy, R., Laske, R. D., Losorelli, S. and Heller, S. (2015). Identification and characterization of mouse otic sensory lineage genes. Front Cell Neurosci 9, 79.

Hatakeyama, M., Opitz, L., Russo, G., Qi, W., Schlapbach, R. and Rehrauer, H. (2016). SUSHI - an exquisite recipe for fully documented, reproducible and reusable NGS data analysis. BMC Bioinformatics 17, 228.

Hidalgo-Sanchez, M., Alvarado-Mallart, R. and Alvarez, I. S. (2000). Pax2, Otx2, Gbx2 and Fgf8 expression in early otic vesicle development. Mech Dev 95, 225–229.

Johnson Chacko, L., Sergi, C., Eberharter, T., Dudas, J., Rask-Andersen, H., Hoermann, R., Fritsch, H., Fischer, N., Glueckert, R. and Schrott-Fischer, A. (2020). Early appearance of key transcription factors influence the spatiotemporal development of the human inner ear. Cell Tissue Res 379, 459–471.

Kaiser, M., Wojahn, I., Rudat, C., Ludtke, T. H., Christoffels, V. M., Moon, A., Kispert, A. and Trowe, M. O. (2021). Regulation of otocyst patterning by Tbx2 and Tbx3 is required for inner ear morphogenesis in the mouse. Development 148.

Kiernan, A. E., Pelling, A. L., Leung, K. K., Tang, A. S., Bell, D. M., Tease, C., Lovell-Badge, R., Steel, K. P. and Cheah, K. S. (2005). Sox2 is required for sensory organ development in the mammalian inner ear. Nature 434, 1031–1035.

Koehler, K. R., Mikosz, A. M., Molosh, A. I., Patel, D. and Hashino, E. (2013). Generation of inner ear sensory epithelia from pluripotent stem cells in 3D culture. Nature 500, 217–221.

Koehler, K. R., Nie, J., Longworth-Mills, E., Liu, X. P., Lee, J., Holt, J. R. and Hashino, E. (2017). Generation of inner ear organoids containing functional hair cells from human pluripotent stem cells. Nat Biotechnol 35, 583–589.

Kolla, L., Kelly, M. C., Mann, Z. F., Anaya-Rocha, A., Ellis, K., Lemons, A., Palermo, A. T., So, K. S., Mays, J. C., Orvis, J., et al. (2020). Characterization of the development of the mouse cochlear epithelium at the single cell level. Nat Commun 11, 2389.

Lavigne-Rebillard, M. and Pujol, R. (1986). Development of the auditory hair cell surface in human fetuses. A scanning electron microscopy study. Anat Embryol (Berl) 174, 369–377.

Lavigne-Rebillard, M. and Pujol, R. (1987). Surface aspects of the developing human organ of Corti. Acta Otolaryngol Suppl 436, 43–50.

Lavigne-Rebillard, M. and Pujol, R. (1988). Hair cell innervation in the fetal human cochlea. Acta Otolaryngol 105, 398–402.

Leger, S. and Brand, M. (2002). Fgf8 and Fgf3 are required for zebrafish ear placode induction, maintenance and inner ear patterning. Mech Dev 119, 91–108.

Li, Y., Liu, H., Giffen, K. P., Chen, L., Beisel, K. W. and He, D. Z. Z. (2018). Transcriptomes of cochlear inner and outer hair cells from adult mice. Sci Data 5, 180199.

Litsiou, A., Hanson, S. and Streit, A. (2005). A balance of FGF, BMP and WNT signalling positions the future placode territory in the head. Development 132, 4051–4062.

Locher, H., de Groot, J. C., van Iperen, L., Huisman, M. A., Frijns, J. H. and Chuva de Sousa Lopes, S. M. (2014). Distribution and development of peripheral glial cells in the human fetal cochlea. Plos One 9, e88066.

Locher, H., de Groot, J. C., van Iperen, L., Huisman, M. A., Frijns, J. H. and Chuva de Sousa Lopes, S. M. (2015). Development of the stria vascularis and potassium regulation in the human fetal cochlea: Insights into hereditary sensorineural hearing loss. Dev Neurobiol 75, 1219–1240.

Locher, H., Frijns, J. H., van Iperen, L., de Groot, J. C., Huisman, M. A. and Chuva de Sousa Lopes, S. M. (2013). Neurosensory development and cell fate determination in the human cochlea. Neural Dev 8, 20.

Luo, X. J., Deng, M., Xie, X., Huang, L., Wang, H., Jiang, L., Liang, G., Hu, F., Tieu, R., Chen, R., et al. (2013). GATA3 controls the specification of prosensory domain and neuronal survival in the mouse cochlea. Hum Mol Genet 22, 3609–3623.

Maroon, H., Walshe, J., Mahmood, R., Kiefer, P., Dickson, C. and Mason, I. (2002). Fgf3 and Fgf8 are required together for formation of the otic placode and vesicle. Development 129, 2099–2108.

Martik, M. L. and Bronner, M. E. (2021). Riding the crest to get a head: neural crest evolution in vertebrates. Nat Rev Neurosci 22, 616–626.

Martin, K. and Groves, A. K. (2006). Competence of cranial ectoderm to respond to Fgf signaling suggests a two-step model of otic placode induction. Development 133, 877–887.

McCarthy DJ, C. K., Lun ATL, Willis QF (2017). Scater: pre-processing, quality control, normalisation and visualisation of single-cell RNA-seq data in R. Bioinformatics 33, 1179–1186.

Mendez-Maldonado, K., Vega-Lopez, G. A., Aybar, M. J. and Velasco, I. (2020). Neurogenesis From Neural Crest Cells: Molecular Mechanisms in the Formation of Cranial Nerves and Ganglia. Front Cell Dev Biol 8, 635.

Menendez, L., Trecek, T., Gopalakrishnan, S., Tao, L., Markowitz, A. L., Yu, H. V., Wang, X., Llamas, J., Huang, C., Lee, J., et al. (2020). Generation of inner ear hair cells by direct lineage conversion of primary somatic cells. Elife 9.

Moore, S. T., Nakamura, T., Nie, J., Solivais, A. J., Aristizábal-Ramírez, I., Ueda, Y., Mayakannan, M., Reddy, V. S., Romano, D. R., Perrin, B. J., et al. (2022). Generating high-fidelity cochlear organoids from human pluripotent stem cells. https://ssrn.com/abstract=4170188 or http://dx.doi.org/10.2139/ssrn.4170188.

Munnamalai, V. and Fekete, D. M. (2017). Building the human inner ear in an organoid. Nat Biotechnol 35, 518–520.

Nie, J. and Hashino, E. (2020). Generation of inner ear organoids from human pluripotent stem cells. Methods Cell Biol 159, 303–321.

Nie, J., Ueda, Y., Solivais, A. J. and Hashino, E. (2022). CHD7 regulates otic lineage specification and hair cell differentiation in human inner ear organoids. Nat Commun 13, 7053.

Nist-Lund, C., Kim, J. and Koehler, K. R. (2022). Advancements in inner ear development, regeneration, and repair through otic organoids. Current Opinion in Genetics & Development 76.

Noda, T., Meas, S. J., Nogami, J., Amemiya, Y., Uchi, R., Ohkawa, Y., Nishimura, K. and Dabdoub, A. (2018). Direct Reprogramming of Spiral Ganglion Non-neuronal Cells into Neurons: Toward Ameliorating Sensorineural Hearing Loss by Gene Therapy. Front Cell Dev Biol 6, 16.

O’Rahilly, R. (1963). The Early Development of the Otic Vesicle in Staged Human Embryos. J. Embryol. exp. Morph 11, 741–755.

Pechriggl, E. J., Bitsche, M., Glueckert, R., Rask-Andersen, H., Blumer, M. J., Schrott-Fischer, A. and Fritsch, H. (2015). Development of the innervation of the human inner ear. Dev Neurobiol 75, 683–702.

Petitpre, C., Faure, L., Uhl, P., Fontanet, P., Filova, I., Pavlinkova, G., Adameyko, I., Hadjab, S. and Lallemend, F. (2022). Single-cell RNA-sequencing analysis of the developing mouse inner ear identifies molecular logic of auditory neuron diversification. Nat Commun 13, 3878.

Pollen, A. A., Bhaduri, A., Andrews, M. G., Nowakowski, T. J., Meyerson, O. S., Mostajo-Radji, M. A., Di Lullo, E., Alvarado, B., Bedolli, M., Dougherty, M. L., et al. (2019). Establishing Cerebral Organoids as Models of Human-Specific Brain Evolution. Cell 176, 743–756 e717.

Pujol, R. and Lavigne-Rebillard, M. (1985). Early stages of innervation and sensory cell differentiation in the human fetal organ of Corti. Acta Otolaryngol Suppl 423, 43–50.

Puligilla, C., Dabdoub, A., Brenowitz, S. D. and Kelley, M. W. (2010). Sox2 induces neuronal formation in the developing mammalian cochlea. J Neurosci 30, 714–722.

Qi, Y., Zhang, X. J., Renier, N., Wu, Z., Atkin, T., Sun, Z., Ozair, M. Z., Tchieu, J., Zimmer, B., Fattahi, F., et al. (2017). Combined small-molecule inhibition accelerates the derivation of functional cortical neurons from human pluripotent stem cells. Nat Biotechnol 35, 154–163.

Radde-Gallwitz, K., Pan, L., Gan, L., Lin, X., Segil, N. and Chen, P. (2004). Expression of Islet1 marks the sensory and neuronal lineages in the mammalian inner ear. J Comp Neurol 477, 412–421.

Ranum, P. T., Goodwin, A. T., Yoshimura, H., Kolbe, D. L., Walls, W. D., Koh, J. Y., He, D. Z. Z. and Smith, R. J. H. (2019). Insights into the Biology of Hearing and Deafness Revealed by Single-Cell RNA Sequencing. Cell Rep 26, 3160–3171 e3163.

Renauld, J. M., Khan, V. and Basch, M. L. (2022). Intermediate Cells of Dual Embryonic Origin Follow a Basal to Apical Gradient of Ingression Into the Lateral Wall of the Cochlea. Front Cell Dev Biol 10, 867153.

Riccomagno, M. M., Takada, S. and Epstein, D. J. (2005). Wnt-dependent regulation of inner ear morphogenesis is balanced by the opposing and supporting roles of Shh. Genes Dev 19, 1612–1623.

Roccio, M. (2021). Directed differentiation and direct reprogramming: Applying stem cell technologies to hearing research. Stem Cells 39, 375–388.

Roccio, M. and Edge, A. S. B. (2019). Inner ear organoids: new tools to understand neurosensory cell development, degeneration and regeneration. Development 146.

Roccio, M., Perny, M., Ealy, M., Widmer, H. R., Heller, S. and Senn, P. (2018). Molecular characterization and prospective isolation of human fetal cochlear hair cell progenitors. Nat Commun 9, 4027.

Saint-Jeannet, J. P. and Moody, S. A. (2014). Establishing the pre-placodal region and breaking it into placodes with distinct identities. Dev Biol 389, 13–27.

Sans, A. and Dechesne, C. (1985). Early development of vestibular receptors in human embryos. An electron microscopic study. Acta Otolaryngol Suppl 423, 51–58.

Scheffer, D. I., Shen, J., Corey, D. P. and Chen, Z. Y. (2015). Gene Expression by Mouse Inner Ear Hair Cells during Development. J Neurosci 35, 6366–6380.

Schindelin, J., Arganda-Carreras, I., Frise, E., Kaynig, V., Longair, M., Pietzsch, T., Preibisch, S., Rueden, C., Saalfeld, S., Schmid, B., et al. (2012). Fiji: an open-source platform for biological-image analysis. Nat Methods 9, 676–682.

Senn, P., Mina, A., Volkenstein, S., Kranebitter, V., Oshima, K. and Heller, S. (2019). Progenitor Cells from the Adult Human Inner Ear. THE ANATOMICAL RECORD.

Shengyang Yu, K., Frumm, S. M., Park, J. S., Lee, K., Wong, D. M., Byrnes, L., Knox, S. M., Sneddon, J. B. and Tward, A. D. Development of the Mouse and Human Cochlea at Single Cell Resolution. https://www.biorxiv.org/content/10.1101/739680v2.

Soldatov, R., Kaucka, M., Kastriti, M. E., Petersen, J., Chontorotzea, T., Englmaier, L., Akkuratova, N., Yang, Y., Haring, M., Dyachuk, V., et al. (2019). Spatiotemporal structure of cell fate decisions in murine neural crest. Science 364.

Steel, K. P. and Barkway, C. (1989). Another role for melanocytes: their importance for normal stria vascularis development in the mammalian inner ear. Development 107, 453–463.

Steevens, A. R., Sookiasian, D. L., Glatzer, J. C. and Kiernan, A. E. (2017). SOX2 is required for inner ear neurogenesis. Sci Rep 7, 4086.

Steinhart, M. R., Serdy, S. A., Van der Valk, W. H., Zhang, J. Y., Kim, J., Lee, J. and Koehler, K. R. Defining inner ear cell type specification at single-cell resolution in a model of human cranial development. Ssrn Electron J 2021,1–58, https://doi.org/10.2139/ssrn.3974124.

Steventon, B., Mayor, R. and Streit, A. (2012). Mutual repression between Gbx2 and Otx2 in sensory placodes reveals a general mechanism for ectodermal patterning. Dev Biol 367, 55–65.

Steventon, B., Mayor, R. and Streit, A. (2014). Neural crest and placode interaction during the development of the cranial sensory system. Dev Biol 389, 28–38.

Stuart, T., Butler, A., Hoffman, P., Hafemeister, C., Papalexi, E., Mauck, W. M., 3rd, Hao, Y., Stoeckius, M., Smibert, P. and Satija, R. (2019). Comprehensive Integration of Single-Cell Data. Cell 177, 1888–1902 e1821.

Sun, Y., Wang, L., Zhu, T., Wu, B., Wang, G., Luo, Z., Li, C., Wei, W. and Liu, Z. (2022). Single-cell transcriptomic landscapes of the otic neuronal lineage at multiple early embryonic ages. Cell Rep 38, 110542.

Taylor, R. R., Jagger, D. J., Saeed, S. R., Axon, P., Donnelly, N., Tysome, J., Moffatt, D., Irving, R., Monksfield, P., Coulson, C., et al. (2015). Characterizing human vestibular sensory epithelia for experimental studies: new hair bundles on old tissue and implications for therapeutic interventions in ageing. Neurobiol Aging 36, 2068–2084.

Uniken Venema, W. T. C., Ramirez-Sanchez, A. D., Bigaeva, E., Withoff, S., Jonkers, I., McIntyre, R. E., Ghouraba, M., Raine, T., Weersma, R. K., Franke, L., et al. (2022). Gut mucosa dissociation protocols influence cell type proportions and single-cell gene expression levels. Sci Rep 12, 9897.

van der Valk, W. H., Steinhart, M. R., Zhang, J. and Koehler, K. R. (2021). Building inner ears: recent advances and future challenges for in vitro organoid systems. Cell Death Differ 28, 24–34.

van der Valk, W. H., van Beelen, E. S. A., Steinhart, M. R., Nist-Lund, C., de Groot, J. C. M. J., van Benthem, P. P. G., Koehler, K. R. and Locher, H. A Single-Cell Level Comparison of Human Inner Ear Organoids and the Human Cochlea and Vestibular Organs. BioRxiv preprint doi: https://doi.org/10.1101/2022.09.28.50983.

Wahle, P., Brancati, G., Harmel, C., He, Z., Gut, G., Santos, A., Yu, Q., Noser, P., Fleck, J. S., Gjeta, B., et al. Multimodal spatiotemporal phenotyping of human organoid development. bioRxiv, DOI: https://doi.org/10.1101/2022.03.16.484396

Waldhaus, J., Durruthy-Durruthy, R. and Heller, S. (2015). Quantitative High-Resolution Cellular Map of the Organ of Corti. Cell Reports 11, 1385–1399.

Wang, T., Ling, A. H., Billings, S. E., Hosseini, D. K., Vaisbuch, Y., Kim, G. S., Atkinson, P. J., Sayyid, Z. N., Aaron, K. A., Wagh, D., et al. Single-cell transcriptomic atlas reveals increased 3 regeneration in diseased human inner ears. BioRxiv preprint doi: https://doi.org/10.1101/2022.10.29.514378.

Wilkerson, B. A., Zebroski, H. L., Finkbeiner, C. R., Chitsazan, A. D., Beach, K. E., Sen, N., Zhang, R. C. and Bermingham-McDonogh, O. (2021). Novel cell types and developmental lineages revealed by single-cell RNA-seq analysis of the mouse crista ampullaris. Elife 10.

Wilson, P. A. and Hemmati-Brivanlou, A. (1995). Induction of epidermis and inhibition of neural fate by Bmp-4. Nature 376, 331–333.

Wiwatpanit, T., Lorenzen, S. M., Cantu, J. A., Foo, C. Z., Hogan, A. K., Marquez, F., Clancy, J. C., Schipma, M. J., Cheatham, M. A., Duggan, A., et al. (2018). Trans-differentiation of outer hair cells into inner hair cells in the absence of INSM1. Nature 563, 691–695.

Wu, D. K. and Kelley, M. W. (2012). Molecular mechanisms of inner ear development. Cold Spring Harb Perspect Biol 4, a008409.

Xu, Y., Zhang, T., Zhou, Q., Hu, M., Qi, Y., Xue, Y., Wang, L., Nie, Y., Bao, Z. and Shi, W. A single-cell transcriptome atlas of human early embryogenesis. bioRxiv preprint doi: https://doi.org/10.1101/2021.11.30.470583.

Yasuda, M., Yamada, S., Uwabe, C., Shiota, K. and Yasuda, Y. (2007). Three-dimensional analysis of inner ear development in human embryos. Anat Sci Int 82, 156–163.

Zelarayan, L. C., Vendrell, V., Alvarez, Y., Dominguez-Frutos, E., Theil, T., Alonso, M. T., Maconochie, M. and Schimmang, T. (2007). Differential requirements for FGF3, FGF8 and FGF10 during inner ear development. Dev Biol 308, 379–391.

